# Phosphatidylserine within the Viral Membrane Enhances Chikungunya Virus Infectivity in a Cell-type Dependent Manner

**DOI:** 10.1101/2022.01.14.476428

**Authors:** Kerri L. Miazgowicz, Judith Mary Reyes Ballista, Marissa D. Acciani, Ariana R. Jimenez, Ryan S. Belloli, Avery M. Duncan, Katherine E. Havranek, Melinda A. Brindley

## Abstract

Chikungunya virus (CHIKV), an alphavirus of the *Togaviridae* family, is the causative agent of the human disease chikungunya fever (CHIKF), which is characterized by debilitating acute and chronic arthralgia. No licensed vaccines or antivirals exist for CHIKV. Preventing the attachment of viral particles to host cells is an attractive intervention strategy. Viral entry of enveloped viruses from diverse families including *Filoviridae* and *Flaviviridae* is mediated or enhanced by phosphatidylserine receptors (PSRs). PSRs facilitate the attachment of enveloped viruses to cells by binding to exposed phosphatidylserine (PS) in the viral lipid membrane - a process termed viral apoptotic mimicry. To investigate the role of viral apoptotic mimicry during CHIKV infection, we produced viral particles with discrete amounts of exposed PS on the virion envelope by exploiting the cellular distribution of phospholipids at the plasma membrane. We found that CHIKV particles containing high outer leaflet PS (produced in cells lacking flippase activity) were more infectious in Vero cells than particles containing low levels of outer leaflet PS (produced in cells lacking scramblase activity). However, the same viral particles were similarly infectious in NIH3T3 and HAP1 cells, suggesting PS levels can influence infectivity only in cells with high levels of PSRs. Interestingly, PS-dependent CHIKV entry was observed in mosquito Aag2 cells, but not C6/36 cells. These data demonstrate that CHIKV entry via viral apoptotic mimicry is cell-type dependent. Furthermore, viral apoptotic mimicry has a mechanistic basis to influence viral dynamics *in vivo* in both the human and mosquito host.

**Importance:** Outbreaks of Chikungunya virus (CHIKV) have occurred throughout Africa, Asia, and Europe. Climate change permits the expansion of *Aedes* mosquito vectors into more temperate regions, broadening the geographic range and increasing the frequency of future human outbreaks. The molecular basis underlying the broad host and cellular tropism of CHIKV remains unresolved. While several host molecules have been implicated in CHIKV viral attachment and entry, the role of lipid-mediated attachment (viral apoptotic mimicry) is unclear. We observed that higher levels of externalized phosphatidylserine (PS) in the viral lipid bilayer correlated with enhanced CHIKV infectivity in mammalian cells abundant with PS receptors and lacking alternative attachment factors. Interestingly, CHIKV infection in mosquito Aag2 cells was also affected by viral PS accessibility. This study further delineates the role of virus-cell attachment molecules in CHIKV infection. Viral apoptotic mimicry has potential to influence CHIKV dynamics *in vivo* in both the human and mosquito host.

## Introduction

Chikungunya virus (CHIKV) is the causative agent of the human disease chikungunya fever (CHIKF). CHIKF develops in approximately 82-95% of infected individuals (1, 2), and is often characterized by debilitating arthralgia in the joints which becomes chronic in 12-36% of cases (3, 4). Other common symptoms include rash, fever, headache and in extreme but rare cases, death (5). Currently, there are no licensed vaccines or antivirals specific to CHIKV. Humans acquire CHIKV from the bite of infected *Aedes aegypti* or *Aedes albopictus* mosquitoes (6, 7). CHIKV outbreaks were originally limited to Africa or Asia (8, 9), however, modern outbreaks introduced CHIKV throughout the Americas and Europe (10–12). The expansion of mosquito vectors (e.g., *Aedes albopictus*) to temperate regions increases the likelihood of future CHIKV outbreaks. Vector control remains the most effective strategy to limit the spread of CHIKV (12–15). Developing interventions that interrupt transmission is essential to mitigating the global health burden from CHIKF.

CHIKV is an *Alphavirus* within the *Togaviridae* family. CHIKV has a positive-sense single-stranded RNA genome that encodes four non-structural proteins (nsP1-4) and six structural proteins (capsid, E1, E2, E3, 6K and TF) (16, 17). CHIKV virions are enveloped, icosahedral particles, studded with 80 glycoprotein spikes comprised of trimeric E1/E2 heterodimers (18, 19). E2 is associated with cellular attachment (16) and E1 is a class II fusion protein that mediates membrane fusion after internalization (16, 20). After capsid uncoating and genome release, genome replication complexes are formed in cytoplasmic invaginations at the plasma membrane (PM), which serve as the site of particle assembly and budding (21).

Virus-cell attachment is an essential step in viral invasion of the host cell. Matrix remodeling associated 8 (MXRA8) (22), glycosaminoglycans (GAGs) such as heparan sulfate (HS) (23–26), C-type lectins including DC-SIGN (27, 28), prohibitin 1 (PHB-1) (29) and phosphatidylserine (PS) receptors such as TIM-1 (30–32) or CD300a (33) have all been implicated in promoting CHIKV entry. The role of MXRA8 in CHIKV pathogenesis has recently been investigated *in vivo* (22, 34). While MXRA8-deficient mice did not develop joint inflammation, infectious virus was still detected in peripheral tissues during acute infection (34), supporting the notion that alternative surface molecules are involved in mediating viral establishment and dissemination. However, none of the identified binding partners are essential to CHIKV infection. Thus, the broad host and cellular tropism of CHIKV may stem from its ability to bind a multitude of molecules present on the cellular surface as opposed to a single ubiquitous factor.

PS on the cell exterior provides a diverse array of physiological functions including cell signaling and membrane fluidity (reviewed in (35)). As such, cells strongly regulate PS orientation within the plasma membrane (PM) to prevent the premature display of PS on the exterior of the cell by restricting PS to the cytosolic leaflet (36). Type 4 P-type ATPases (P4-ATPases), termed flippases, actively translocate PS from the exoplasmic leaflet of the lipid bilayer to the cytosolic leaflet in an ATP-dependent manner to maintain an asymmetric PS gradient in healthy cells (37, 38). P4-ATPases require a subunit from the CDC50 family to promote appropriate cellular localization and flippase activity (39–41). Apoptotic induction results in the irreversible inactivation of P4-ATPases through caspase cleavage (42), and activation of a second class of phospholipid regulatory enzymes, termed scramblases (43). Scramblases indiscriminately shuffle phospholipids between the inner and outer leaflets (43). Some scramblases, including transmembrane protein 16F (TMEM16F), can undergo a reversible activation through calcium signaling (44), while others, such as XK-related protein 8 (XKR8), are irreversibly activated from caspase cleavage after apoptotic initiation (43).

Phosphatidylserine receptors (PSRs) can facilitate pathogen attachment to cells (30, 31, 45–48). The induction of cellular apoptosis after viral infection was traditionally considered a host-driven antiviral response. However, mounting evidence from several viral families including *Filoviridae* (e.g. Ebola virus (31, 49)) and *Flaviviridae* (e.g. Dengue virus (30, 33, 50)) illustrates that virions budding from an apoptotic cell can confer pro-viral effects. Viruses containing outer-leaflet associated PS in the viral envelope can engage PSRs on host cells, mimicking apoptotic bodies and triggering internalization (51). As CHIKV buds from the PM of an infected cell, PS externalization during apoptosis could enhance the ability of virions to attach to nearby uninfected cells.

In this study, we exploited PM-associated flippases and scramblases to modify the natural phospholipid dynamics within the lipid bilayer of the cellular PM to produce CHIKV virions with distinct levels of external PS (low, moderate, or high). We used these particles to assess the role of viral apoptotic mimicry during CHIKV infection. We postulated that increased PS levels in the outer leaflet of the virion envelope would promote cellular attachment, resulting in a more efficient infection in cells containing PSRs. Understanding the variation in entry efficiency among CHIKV attachment factors broadens our understanding of the molecular basis for the diverse species and tissue tropism of CHIKV. Viral dynamics and the cross-species transmission of CHIKV between mammalian and mosquito hosts are likely influenced by the assortment of cellular attachment factors across cell types.

## Results

### TIM-1 enhances CHIKV infection in a cell-dependent manner

Previous studies demonstrated that CHIKV infection could be enhanced by the addition of entry factors including MXRA8, lectin binding proteins, and phosphatidylserine receptors (31, 32, 45). First, we sought to confirm previous findings in 293T cells and evaluate if an infection enhancement is observed in commonly used cell lines including HAP1 and Vero cells. Cells were transfected with a plasmid encoding hTIM-1 fused with GFP (hTIM-1-GFP), MXRA8, L-SIGN or a control GFP plasmid. Production of exogenous hTIM-1-GFP, MXRA8, or L-SIGN in 293T cells, which do not natively produce these proteins (22, 31, 52), was verified by flow cytometry (**Figure 1A**). We then assessed transfected cells (GFP^+^) for CHIKV infection (mKate^+^) in comparison to GFP-only control wells. Corroborating previous studies (31, 32, 45), the production of hTIM-1-GFP resulted in a 4-fold increase in CHIKV infection in 293T cells relative to GFP-only transfected cells (**Figure 1B**). However, production of exogenous TIM-1-GFP did not increase CHIKV entry into HAP1 or Vero-hSLAM (VeroS) cells (**Figure 1C, D**). CHIKV more readily infected 293T and HAP cells producing MXRA8 (**Figure 1B, C**). Yet, production of MXRA8 did not enhance CHIKV entry into VeroS cells (**Figure 1D)**. L-SIGN addition resulted in a 5-fold increase in CHIKV infection in 293T cells but had no effect on HAP1 or VeroS cells (**Figure 1B-D**). Overproduction of individual CHIKV attachment factors facilitated CHIKV infection in a cell-type dependent manner.

**Figure 1.**
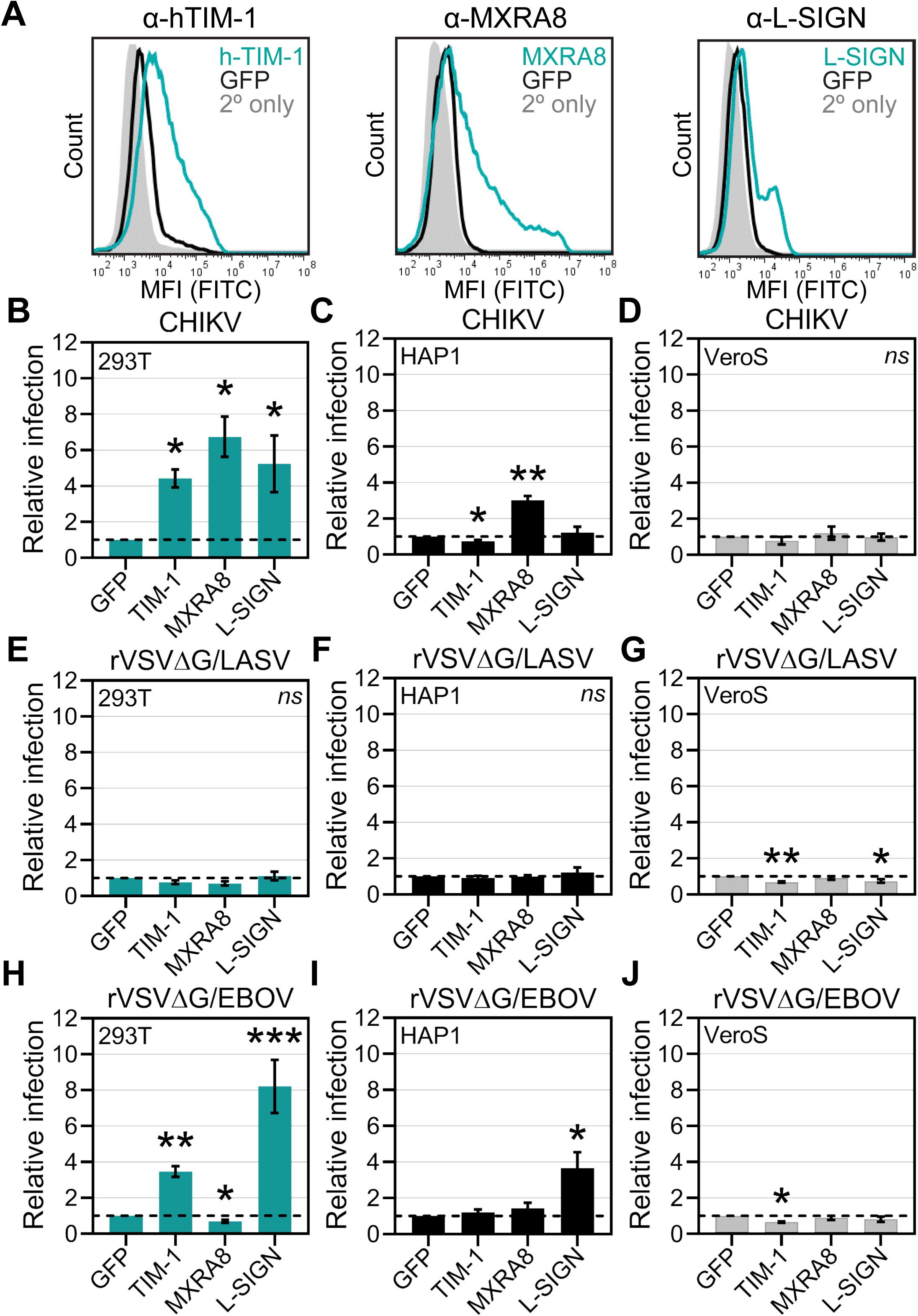
Overexpression of TIM-1 enhances CHIKV infection in a cell-dependent manner. (A) 293T cells were assessed for the surface presentation of known CHIKV attachment factors (TIM-1, MXRA8, or L-SIGN) via flow cytometry. 293T cells were transfected with either TIM-1, MXRA9, L-SIGN, or GFP 24 hrs prior to the addition of primary antibodies specific against hTIM-1, MXRA8, or L-SIGN. 24 hrs post transfection, 293T, HAP1, and VeroS cells were inoculated with either (**B-D**) mKate-expressing CHIKV strain 181/c25, (**E-G**) recombinant vesicular stomatitis virus containing the Lassa virus glycoprotein (rVSVΔG/LASV), or (**H-J**) rVSV studded with the Ebola virus glycoprotein (rVSVΔG/EBOV) for 1 hr. 12 hrs post infection, cells were assessed for CHIKV infection (mKate^+^) and transfection efficiency (GFP^+^) via flow cytometry. Relative infection was calculated as the proportion of cells infected with CHIKV (mKate^+^) among transfected cells (GFP^+^) normalized to infection levels in a GFP only control well. At least three independent replicates were performed with each bar representing the mean and error (±SEM) with an unpaired parametric Student’s T-test with unequal variance (Welch’s correction) was used to determine statistical significance, where * (*p* < 0.05), ** (*p* < 0.01), *** (*p* < 0.001).

To confirm that the infection enhancements were specific to CHIKV, we infected cells producing exogenous entry factors with recombinant vesicular stomatitis virus (rVSV) containing the Lassa virus glycoprotein (rVSVΔG/LASV) (52). Both 293T and HAP1 cells produce properly glycosylated alpha-dystroglycan, the high affinity receptor for Lassa virus (53, 54), whereas VeroS cells do not (47, 55). As expected, the overproduction of TIM-1, MXRA8, or L-SIGN did not significantly affect the entry of rVSVΔG/LASV into either 293T, HAP1, or VeroS cells (**Figure 1E-G**).

Lastly, we used rVSV particles containing the Ebola virus glycoprotein (rVSVΔG/EBOV), which has previously demonstrated a viral enhancement with phosphatidylserine receptors, but not with CHIKV-specific receptor MXRA8. Entry of rVSVΔG/EBOV particles (49) was enhanced in 293T cells producing TIM-1 (**Figure 1H**). As expected, MXRA8 did not increase rVSVΔG/EBOV infection in 293T, HAP1, or VeroS cells (**Figure 1H-J**). L-SIGN enhanced rVSVΔG/EBOV infection by 8-fold in 293T cells and by 3.5- fold in HAP1 cells but had no effect in VeroS cells (**Figure 1H-J**). Together, these data suggest that the role of viral apoptotic mimicry in infection is cell-type specific and depends on the receptors and attachment factors present on the cellular surface.

### Flippase and scramblase knockout cells alter natural PS externalization

To further evaluate the role of PS and PSRs in CHIKV infection, we used cells with modified PS translocation dynamics at the plasma membrane (PM) to generate CHIKV virions with discrete levels of exposed PS on the viral envelope. Knocking out (KO) CDC50A in HAP1 cells (HAP1ΔCDC50A) eliminates P4-ATPase flippase activity, theoretically resulting in cells with relatively high PS levels in the outer leaflet of the PM. In contrast, deleting XKR8 in HAP1 cells (HAP1ΔXKR8) prevents apoptosis-induced scramblase activity, theoretically resulting in cells with outer leaflets that remain low in PS even during apoptosis.

Flippase and scramblase knockout lines were functionally validated by assessing PS externalization and PM integrity in live cells over time. As expected, knocking out CDC50A resulted in increased basal levels of external PS relative to the parental HAP1 line throughout the time course (**Figure 2A**). Conversely, HAP1ΔXKR8 cells maintained the lowest levels of external PS (**Figure 2A**). The activity of cellular flippases and scramblases is dynamically regulated by stimuli such as apoptosis and calcium influx. To further validate the functional phenotype of our KO lines, we infected cells with CHIKV (strain 181/c25), a known inducer of apoptosis (56). Congruent with basal conditions, apoptotic HAP1ΔCDC50A cells displayed the highest levels of PS in the outer leaflet of the PM and HAP1ΔXKR8 cells displayed the lowest (**Figure 2B**). As hypothesized, CHIKV infection resulted in stronger induction of apoptosis compared to basal levels, while cells lacking XKR8 remained low in outer leaflet PS under both treatments (**Figure 2B**).

**Figure 2.**
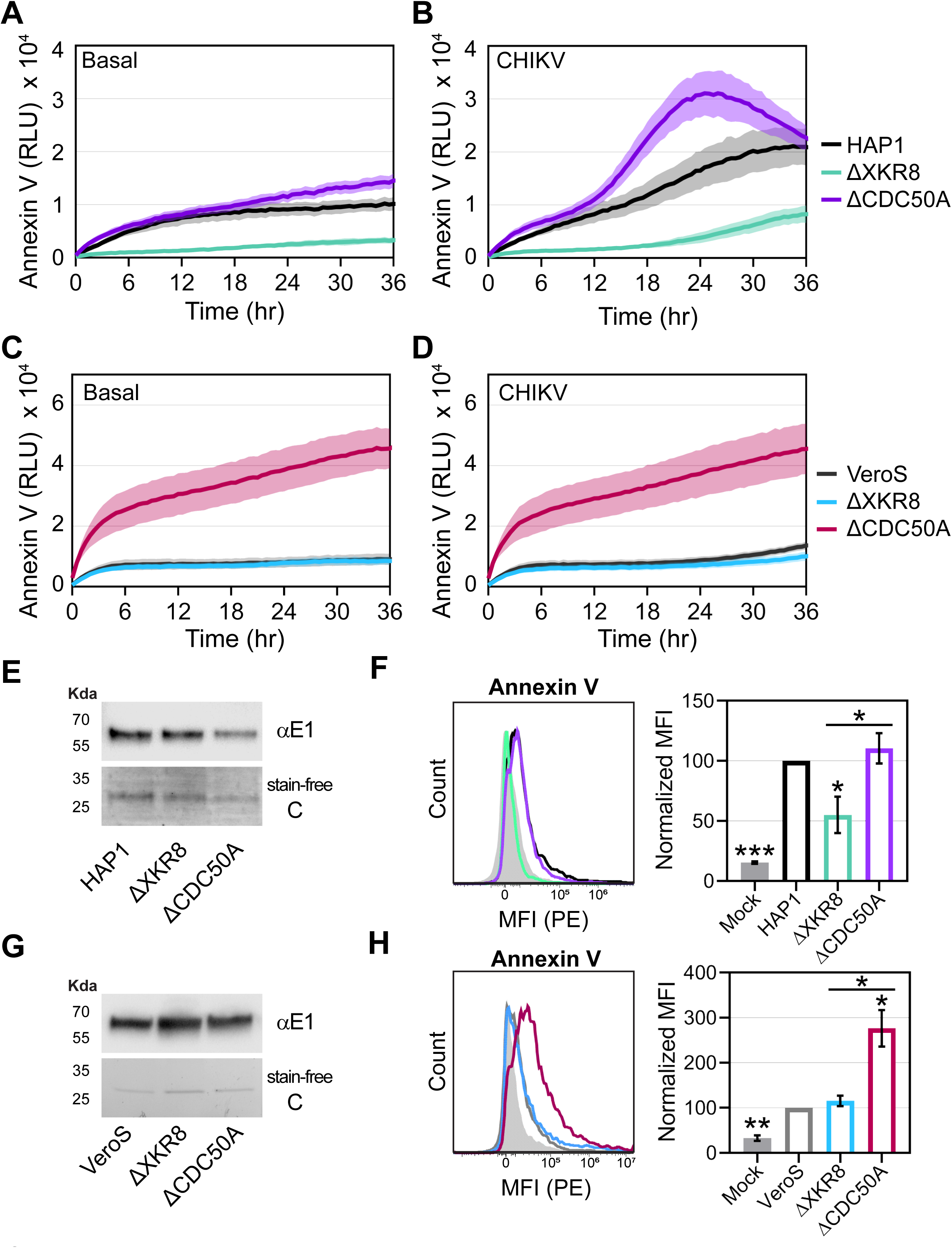
Knocking out flippases and scramblases affects the amount of externalized cellular PS. HAP1 cell lines and VeroS cell lines were monitored for Annexin V binding (RLU) for 36 hrs with automated measurements taken every 30 mins using a GloMax Explorer multimode microplate reader held at 37°C. Parental, scramblase XKR8 KO, and flippase subunit CDC50A KO cells were either (**A**, **C**) untreated (Basal) or treated with (**B**, **D**) viral infection (CHIKV strain 181/c25, MOI 1.0). Three independent replicates were performed with the solid line representing the mean and faded region indicating the error (±SEM). To quantify levels of externalized PS on the CHIKV viral particle, CHIKV was propagated through both HAP1 and VeroS cell lines. (**E, G**) Viral inputs were immunoblotted with an α-CHIKV antibody and accessed for purity using a stain-free gel. (**F, H**) Annexin V conjugated to PE was used to stain normalized amounts of virus-bound beads and quantified via FACs analysis. A bead only control (mock) was used to establish a baseline signal. MFI values from three independent trials were normalized to parental values (HAP1 or VeroS) with the mean and ±SEM displayed. An unpaired parametric T-test with Welch’s correction was used test for statistical significance, where * (*p* < 0.05), ** (*p* < 0.01), *** (*p* < 0.001).

To validate our findings in an additional cell line, both XKR8 and CDC50A were KO in VeroS cells (VeroSΔXKR8 or VeroSΔCDC50A). Consistent with the HAP1 background, VeroSΔCDC50A cells displayed the highest levels of outer leaflet PS in both untreated and CHIKV infected cells (**Figure 2C, D**). In contrast, parental VeroS cells displayed indistinguishable basal levels of externalized PS relative to VeroSΔXKR8 cells (**Figure 2C**). However, all VeroS lines lacked a strong signal of scrambling activity following CHIKV infection (**Figure 2D**).

Next, we assessed the accessibility of PS on the viral envelope to verify that the lipid orientation from the cellular PM, the site of CHIKV budding, is maintained. CHIKV particles collected from each cell line were purified via ultracentrifugation. Normalized genome equivalents were bound to beads for Annexin V immunofluorescent staining and a portion was analyzed for CHIKV protein content. CHIKV glycoprotein (E) was detected at mostly similar intensities from input virus via immunoblotting with α-CHIKV antibody, except that the input from HAP1ΔCDC50A appeared slightly lower (**Figure 2E**). Stain-free protein imaging confirmed the level of capsid (C) protein was similar among the samples (**Figure 2E**). Congruent with PS orientation on the cellular membrane, CHIKV particles produced from CDC50A KO HAP1 cells bound a statistically higher number of annexin V molecules than HAP1ΔXKR8 produced particles (**Figure 2F**). While external PS levels on CHIKV particles produced from XKR8 scramblase KO HAP1 cells were significantly lower than WT, the trend between WT and HAP1ΔCDC50A did not achieve statistical significance, potentially due to a slightly lower amount of HAP1ΔCDC50A input virus (**Figure 2F**). In VeroS cells, similar intensities of CHIKV E were detected via immunoblotting (**Figure 2G**) and C using the stain-free protein staining (**Figure 2G**). Externalized PS detected on CHIKV particles from VeroS cells were consistent with the PS orientation at the PM, where VeroSΔCDC50A particles bound well to annexin V, while particles produced in VeroS was indistinguishable from scramblase KO VeroSΔXKR8 (**Figure 2H**).

### Deleting CDC50A results in higher CHIKV titers in Vero cells

Next, we evaluated if altered localization of cellular PS affected CHIKV viral titers in a multi-cycle replication assay. If apoptotic mimicry enhances CHIKV infection, we expected viral titers to be highest in our flippase KO lines as the PS-high virions produced from these cells should bind PS receptors (PSRs) on naive host cells with increased efficiency in subsequent rounds of infection and boost CHIKV entry. Conversely, the lack of exposed PS on virions produced in scramblase KO cells should limit virion attachment to PSRs, resulting in fewer cells infected and a net decrease in viral titers relative to parental lines.

Contrary to our expectations, CHIKV replication kinetics were similar among HAP1ΔCDC50A, HAP1, and HAP1ΔXKR8 cells when titrated on HAP1 cells (**Figure 3A**). HAP1 cells lack robust PSR expression (Horizon Discovery), and therefore we hypothesized virion infectivity may appear equivalent if attachment occurs through non-apoptotic mimicry associated molecules. Next, we titrated the same viral supernatants on VeroS cells to compare viral titers in cells that contain the PSRs TIM-1 and AXL (47). When titrated on VeroS cells, HAP1ΔCDC50A cells consistently produced higher viral titers than the other HAP1 cell lines (**Figure 3B**). No difference in viral titers was observed between HAP1 and scramblase KO HAP1ΔXKR8 cells (**Figure 3A, B**) despite XKR8 KO cells having substantially lower external PS on the PM throughout the course of CHIKV infection (**Figure 2B**).

**Figure 3.**
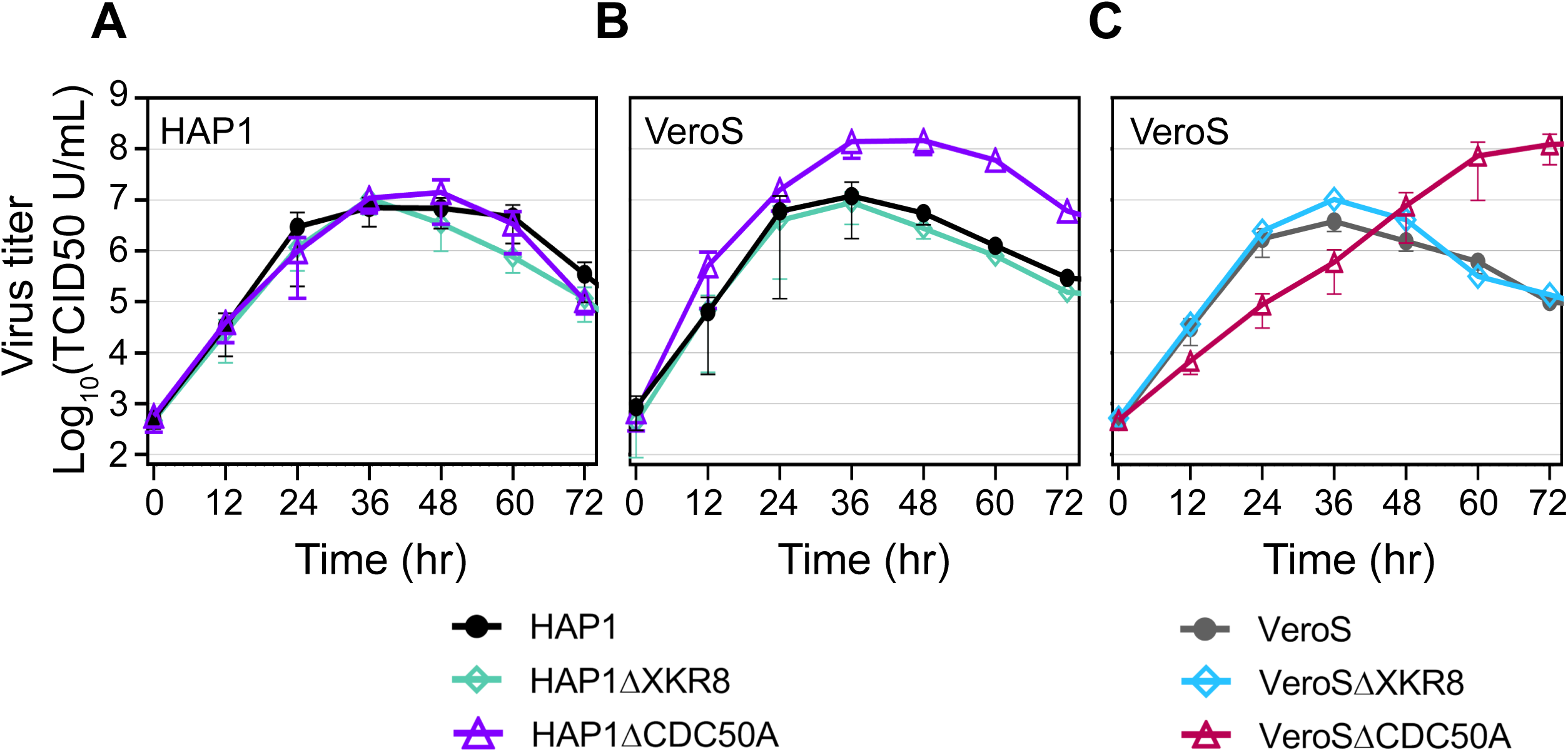
Multi-step CHIKV replication kinetics. Multi-step replication kinetics of CHIKV strain 181/c25 (MOI = 0.01) in parental, scramblase XKR8 KO, or flippase subunit CDC50A KO cells. Cellular supernatants from HAP1 cell lines were titrated on either (**A**) HAP1 cells or (**B**) VeroS cells. (**C**) Cellular supernatants from VeroS cell lines were titrated on VeroS cells. Three independent replicates were performed with error bars representing ±SEM.

Multi-cycle replication assays were also performed in the VeroS background. Interestingly, viral titers from our flippase KO VeroSΔCDC50A cells were initially lower than titers from parental cells, yet surpassed WT levels late in the course of infection (**Figure 3C**). As with the HAP1 background, we found CHIKV replication kinetics were unaffected in our scramblase KO line (VeroSΔXKR8) compared to parental cells (**Figure 3C**). Thus, these data suggest that CHIKV can achieve higher viral titers in the absence of CDC50A when infectivity is quantified on Vero cells, but not HAP1 cells. Further, the lack of accessible PS on the cell surface or the viral particle did not inhibit CHIKV infection relative to wild-type.

### CDC50A KO cells achieve higher viral titers without an increase in entry or cellular spread

Differences in cell permissivity or susceptibility could result in more virions produced per infected cell. This could explain the higher viral titers achieved in a multi-step replication curve from CDC50A KO cells as opposed to increased particle infectivity mediated by improved PS-PSR interactions. To assess for differences in permissivity or susceptibility, we determined the number of particles produced from each cell line by quantifying the number of viral genomes in the cellular supernatant at peak infection (36 hpi) during the multi-cycle replication curve shown in **Figure 3**. Genome equivalents did not significantly vary between HAP1ΔCDC50A, HAP1, or HAP1ΔXKR8 cells (**Figure 4A**). Consistent with the trends observed in the multi-step growth curve (**Figure 3C**), genomes equivalents at 36 hpi were similar from VeroS and VeroSΔXKR8 cells compared to a statistically significant reduction in genomes equivalents from VeroSΔCDC50A cells (**Figure 4B**).

**Figure 4.**
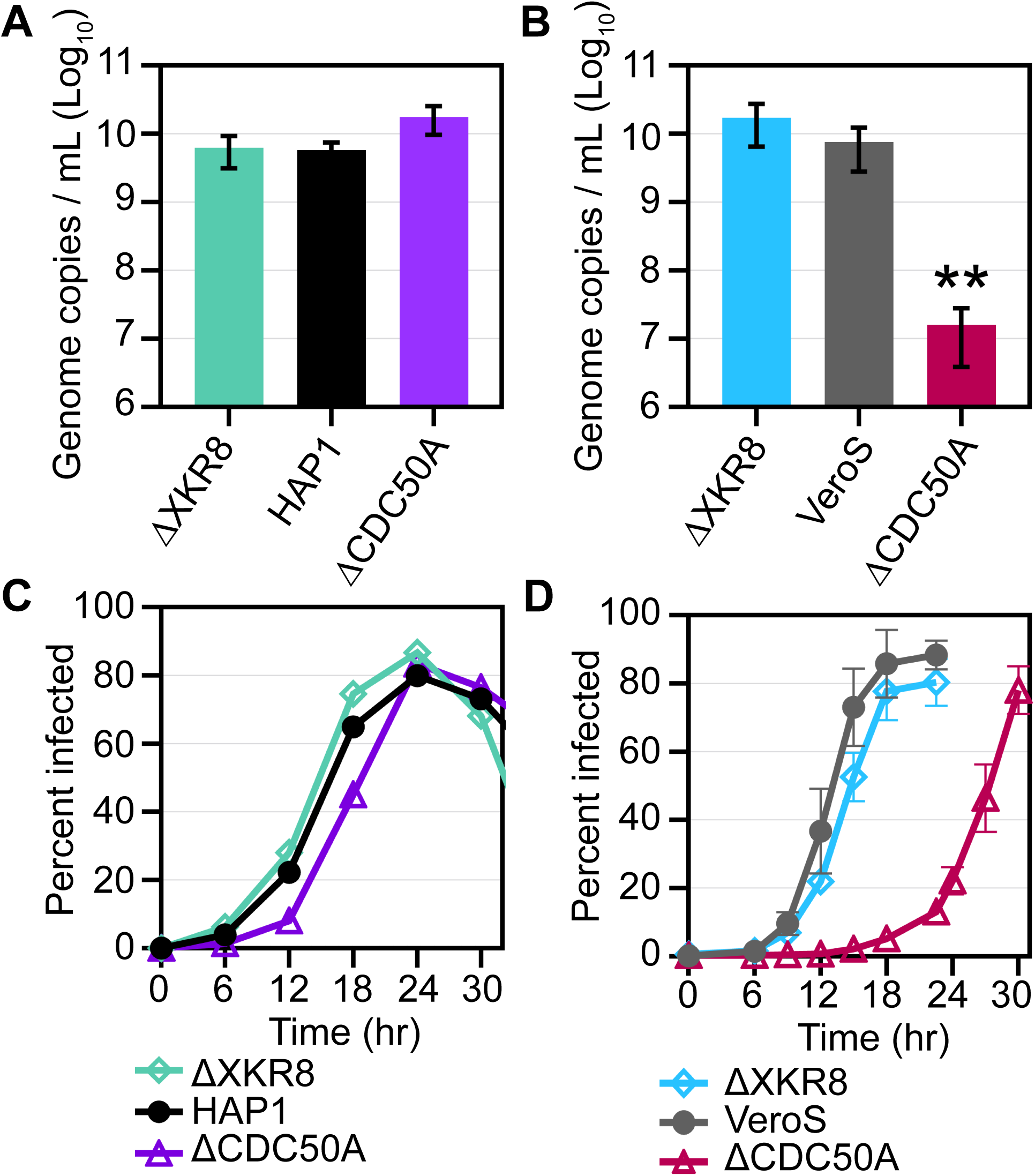
Increased CHIKV titers in CDC50A KO cells are not due to increased viral production or cellular spread. The number of genome copy equivalents per mL were determined from the cellular supernatants collected at 36 hrs post infection (peak infection) from the multi-step replication curve (MOI = 0.01) in Figure 3 with real-time qPCR for each (**A**) HAP1 cell line and (**B**) VeroS cell line. Genome copy equivalents per mL were natural log (ln) transformed prior to performing an unpaired parametric student T-test relative to parental cells (HAP1 or VeroS). CHIKV cellular spread kinetics were quantified by FACs analysis of the percent of GFP^+^ cells over time after CHIKV-*gfp* infection (MOI = 0.1) in (**C**) HAP1 lines or (**D**) VeroS lines. Three independent replicates were conducted with error bars representing ±SEM. Values of significance are shown as * (*p* < 0.05), ** (*p* < 0.01), *** (*p* < 0.001).

We next monitored the number of infected cells over time to discern if CHIKV was spreading through CDC50A KO cells faster than parental or scramblase KO cell lines, which we would hypothesize if PS-PSR interactions contribute to CHIKV infectivity. Contrary to our expectations, all HAP1 cell lines were infected at similar rates across several rounds of CHIKV infection (**Figure 4C**). While CHIKV spread through VeroS and VeroSΔXKR8 cells at similar rates, we observed a delay in the kinetics of CHIKV^+^ VeroSΔCDC50A cells (**Figure 4D**). However, once approximately 10% of the VeroSΔCDC50A population was infected, CHIKV spread at a rate similar to that of VeroS and VeroSΔXKR8 cells (**Figure 4D**). Altering PS orientation at the cellular PM did not alter CHIKV replication kinetics except in VeroSΔCDC50A cells, where viral spread was delayed. Thus, differences in cell permissibility or susceptibility do not explain the overall increased viral titers observed in CDC50A KO cells when titrated on a cell line that contains PSRs.

### CHIKV entry in Vero cells predominately utilizes PS receptors

We hypothesized that virions with increased levels of PS on their outer leaflet (e.g. virus produced in CDC50A KO cells) are more infectious due to a higher propensity to bind to cellular surface PSRs. However, virus produced in cells high in outer leaflet PS have delayed viral spread in VeroSΔCDC50A cells and initially lower titers during in a multi-cycle replication curve when quantified on VeroS cells (**Figure 3C**). We sought to examine why CHIKV infection was delayed in VeroSΔCDC50A cells, but not HAP1ΔCDC50A cells.

First, we performed a high MOI entry experiment and monitored reporter gene expression 12 hours following CHIKV infection in our CDC50A KO, XKR8 KO, and parental cell lines. To ensure we captured a single round of replication, cells were treated with ammonium chloride (NH_4_Cl) either with viral inoculum or 2 hours after infection to prevent endosomal acidification and synchronize infection. As expected, CHIKV infection was blocked when NH_4_Cl was added directly with the inoculum (**Figure 5A, B**). We found that the altered cellular PS levels on our flippase and scramblase KO HAP1 lines did not affect the first round of CHIKV viral entry (**Figure 5A**). In contrast, we observed a 94% reduction in CHIKV infected flippase KO VeroSΔCDC50A cells relative to parental cells in the first round of infection and no reduction in CHIKV^+^ VeroSΔXKR8 cells (**Figure 5B**). This strong entry inhibition corresponds with the low CHIKV titers observed in VeroSΔCDC50A cells early in the multi-step replication curve (**Figure 3C**) and initial delay in viral spread (**Figure 4D**).

**Figure 5.**
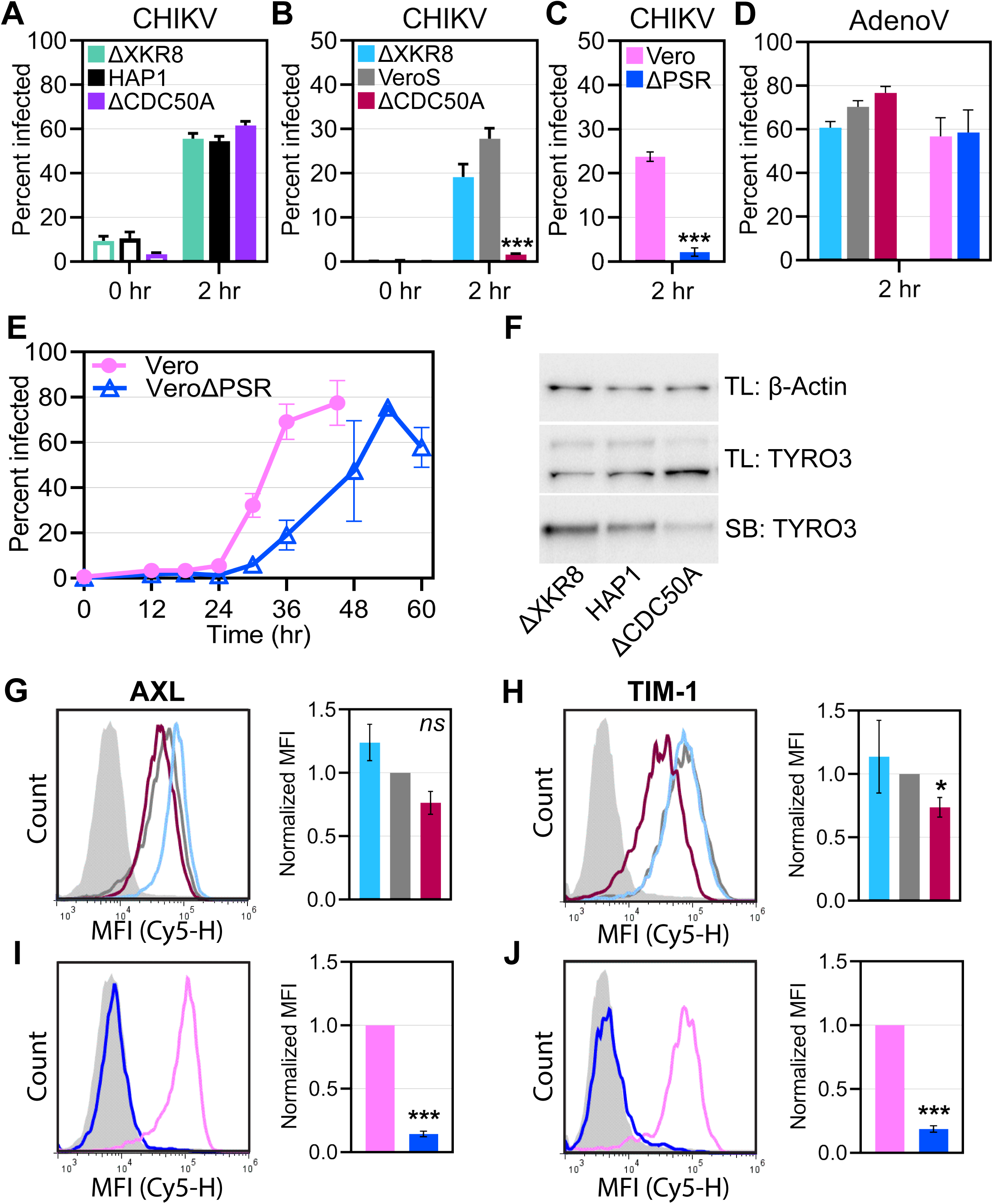
CHIKV entry in Vero cells predominately utilizes PS receptors. CHIKV-*gfp* (strain 181/c25) stock virus was used to infect either (**A**) HAP1 lines (MOI = 10), (**B**) VeroS lines (MOI = 100) or (**C**) Vero lines (MOI =100). 30mM ammonium chloride (NH_4_Cl) was added with the viral inoculum (0hr: hollow bars) or 2 hrs post inoculum removal (2hr: solid bars). At 12 hrs post infection, the percent of cells infected (GFP^+^) was determined by FACs analysis. Three independent replicates were conducted with error bars representing ±SEM. Unpaired parametric student T-tests were performed relative to the parental cell line (HAP1, VeroS, or Vero). (**D**) A viral inoculum was used to achieve ∼60% of cells infected with AdenoV in parental VeroS and Vero cells. Viral inoculum was removed and replaced with complete media 2 hrs post infection. At 20 hrs post infection, the percent of infected cells (GFP^+^) was determined by FACs analysis (**E**) CHIKV cellular spread kinetics were quantified by FACs analysis of the percent of GFP^+^ cells over time after CHIKV-*gfp* infection (MOI = 0.1) in Vero and VeroΔPSR cells. Three independent replicates were conducted with error bars representing ±SEM. (**F**) HAP1, HAP1ΔXKR8, and HAP1ΔCDC50A cells were immunoblotted for the PSR TYRO3 in both the total cell lysate (TB) and surface (SB). β-Actin immunoblotting of the TL served as a protein loading control. A representative image of two independent trials is displayed. The surface presentation of the PSR (**G**) AXL or (**H**) TIM-1 on each VeroS cell line was assessed with immunofluorescence. The mean fluorescence intensity (MFI) was normalized to the parental value within each trial with the mean and ±SEM represented. An unpaired parametric student T-test with a Welch’s correction was performed with mean normalized MFI values relative to parental cells. Surface levels of (**I**) AXL or (**J**) TIM-1 were assessed for Vero and VeroΔPSR cells as described above. Values of significance are shown as * (*p* < 0.05), ** (*p* < 0.01), *** (*p* < 0.001).

Next, we investigated the mechanism behind the observed CHIKV entry inhibition in the flippase KO VeroSΔCDC50A cells. Both the HAP1 and VeroS cell lines lacking CDC50A contain high levels of PS in their outer leaflets (**Figure 2 A-D**). We hypothesized PS may interact with PSR on neighboring cells causing PSR down regulation, inducing a lower steady-state level of PSRs at the plasma membrane. The decreased levels of PSRs on CDC50A KO cells could result in decreased virus entry mediated by viral apoptotic mimicry.

To determine if the CHIKV entry defect in VeroSΔCDC50A cells was associated with a decrease of surface PSRs or an off-target effect of CDC50A deletion, virus entry was assessed in a previously characterized AXL/TIM-1 double knockout Vero cell line (VeroΔPSR) (47) and its parental line (Vero). The absence of TIM-1 and AXL on the cellular surface of VeroΔPSR cells recapitulated the entry defect observed in VeroSΔCDC50A cells (**Figure 5B, C**). In addition, adenovirus infection (non-enveloped, clathrin-mediated entry mechanism) did not result in an entry defect into either VeroSΔCDC50A or VeroΔPSR cells (**Figure 5D**). Lastly, the spread of CHIKV through VeroΔPSR cells was delayed relative to the parental Vero line (**Figure 5E**); reflective of the viral spread kinetics observed among VeroSΔCDC50A cells (**Figure 4D**). Collectively, these data suggest that CHIKV entry defect into VeroSΔCDC50A cells is related to PSRs interactions and not a global viral entry defect.

To investigate if PSRs surface down-regulation was occurring in CDC50A KO cells, the level of PSRs present on the cell surface was compared. According to Horizon Discovery mRNA expression data, HAP1 cells produce the transcripts for the PSR TYRO3, but not other major TIM or TAM family members. Proteins present at the cell surface of HAP1 cells were labeled with biotin, purified, and TYRO3 was detected by immunoblot. Surface TYRO3 levels (SB) were weaker from HAP1ΔCDC50A relative to either parental HAP1 or HAP1ΔXKR8 cells, while total cell lysates (TL) for both β-Actin and TYRO3 was comparable across the three cell lines (**Figure 5F**).

VeroS and Vero cells produce both TIM-1 and AXL (47), but not TYRO3 or Mer (57). Immunofluorescence staining indicates that VeroSΔCDC50A cells display lower levels of both AXL (**Figure 5G**) and TIM-1 (**Figure 5H**) compared to VeroS and VeroSΔXKR8 cells. Further, VeroΔPSR cells were confirmed to lack the surface presentation of both AXL (**Figure 5I**) and TIM-1 (**Figure 5J**).

These data support that PSR surface downregulation is associated with the CHIKV entry defect in flippase KO VeroSΔCDC50A cells. Interestingly, while PSR downregulation also occurs in HAP1ΔCDC50A cells, CHIKV entry remains unaffected. Thus, these data also suggest that CHIKV entry into Vero cells is dependent on PS-PSR interactions, while entry into HAP1 cells is facilitated via alternative binding partners and is independent of PS-PSR interactions.

### CHIKV infection is enhanced through viral apoptotic mimicry in a cell-type dependent manner in both mammalian and mosquito cells

We infected a panel of commonly used mammalian and insect cell lines with CHIKV virions containing discrete levels of envelope outer leaflet PS to determine the relevance of PS-mediated cellular attachment amidst alternative attachment factors and receptors. Genome equivalents were calculated for each viral inoculum and compared to the tissue culture infectious dose 50 value (TCID_50_) to calculate the particle to TCID_50_ ratio as a metric of particle infectivity on each cell type. We observed a correlation between the levels of PS on the particle and particle infectivity when infecting Vero and VeroS cells (**Figure 6A, B**). Particles produced in HAP1, flippase and scramblase KO cells displayed three levels of PS (**Figure 2F**) and three levels of infectivity into Vero and VeroS cells (**Figure 6A, B**). Whereas virus made in VeroS, flippase and scramblase KO cells only displayed two levels of PS on the virus (**Figure 2H**) and displayed two levels of infectivity (**Figure 6A, B**). Infectivity did not correlate with particle PS when particles were added to either HAP1, VeroΔPSR, or NIH3T3 cells (**Figure 6A, B**). Our data suggest CHIKV entry into Vero cells lines is altered by particle PS levels, while the infectivity of the other tested mammalian cell lines is unaffected.

**Figure 6.**
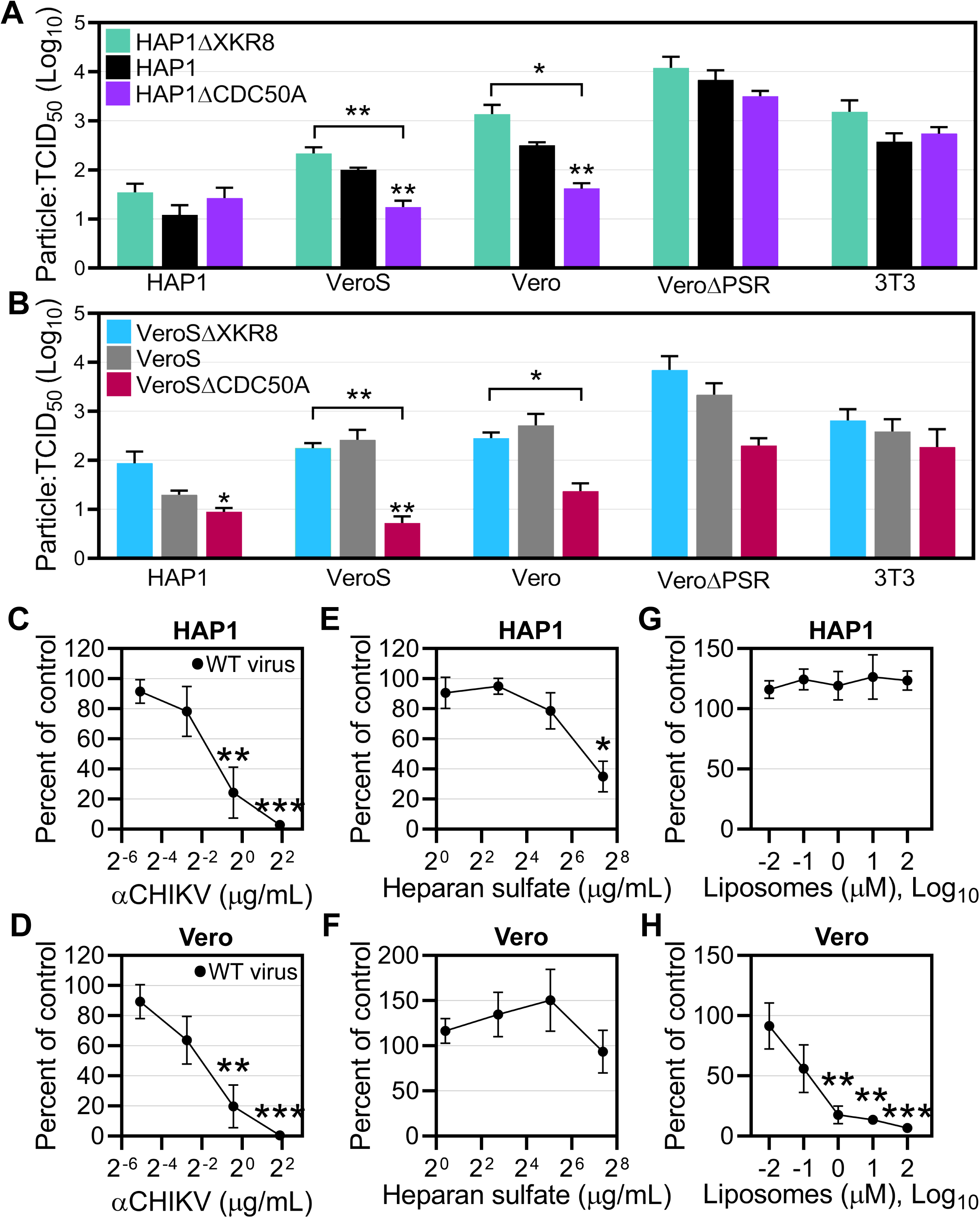
CHIKV viral apoptotic mimicry is cell-type dependent in mammalian cells. The particle (genome copy equivalents) per mL to TCID_50_ units per mL ratio for each sample was used to assess the infectivity of particles produced from (**A**) HAP1 cell lines or (**B**) VeroS cell lines on a panel of commonly used mammalian cell types (human HAP1, monkey Vero, and mouse NIH3T3). CHIKV virions were collected from the producer cell at 24 hrs post CHIKV infection (MOI of 1.0). Congruent with TCID50 methods, equal volumes of collected virus were added to each cell line in the panel without inoculum removal and scored based on CPE and GFP^+^. At least three independent replicates were conducted with bars representing the mean and error (±SEM). Infectivity values were natural log (ln) transformed prior to performing an unpaired parametric student T-test relative to parental cells. (**C-H**) CHIKV-N*luc* stocks were used to assess the ability of (**C, D**) CHIKV antibody, (**E, F**) heparan sulfate, or (**G, H**) PS-containing liposomes (30% PS: 69% PE: 1% PC), to block infections into either HAP1 or Vero mammalian cells at the indicated concentrations. Twenty-four hours following infection the cells were lysed with NanoGlo substrate and lysates were quantified with a GloMax Explorer. At least three independent replicates were conducted with dots representing means and error bars representing ±SEM. Values of significance are shown as * (*p* < 0.05), ** (*p* < 0.01), *** (*p* < 0.001).

Based on the known CHIKV attachment factors present on HAP1 and VeroS cells, we performed competitive inhibition assays to confirm the relevance of glycosaminoglycans and phosphatidylserine receptors in facilitating CHIKV infection. As expected, luciferase signal produced by CHIKV-*Nluc* infection decreased with the addition of increasing concentrations of α-CHIKV antibody in a dose-dependent manner in HAP1 (**Figure 6C**) and Vero (**Figure 6D**) cells. The addition of high concentrations of soluble heparan sulfate competed for CHIKV infection in HAP1 cells (**Figure 6E**) but not Vero cells (**Figure 6F**). CHIKV infection in HAP1 cells was unaffected by the addition of PS containing liposomes (**Figure 6G**). Conversely, Vero cells exhibited dose-dependent inhibition, where a ∼90% reduction in infection was achieved with 100 µM liposomes (**Figure 6H**). These data support the cell-type dependence of CHIKV entry factors, including viral apoptotic mimicry in specific mammalian cells.

The attachment factors promoting CHIKV entry into insect cells remain undefined. Relative to mammalian cells, mosquito cells contain high levels of phosphatidylethanolamine (PE) (58–60), which is another negatively charged phospholipid implicated in binding to PSRs and viral apoptotic mimicry (48). Viral particles produced in the HAP1 cell lines did not alter infectivity in C6/36 cells (**Figure 7A**), but HAP1ΔCDC50A derived particles, high in outer leaflet PS, increased particle infectivity in Aag2 cells (**Figure 7B**). In contrast, virus produced from flippase activity KO VeroS cells enhanced CHIKV infectivity in both mosquito C6/36 (**Figure 7C)** and Aag2 **(Figure 7D**) cells.

**Figure 7.**
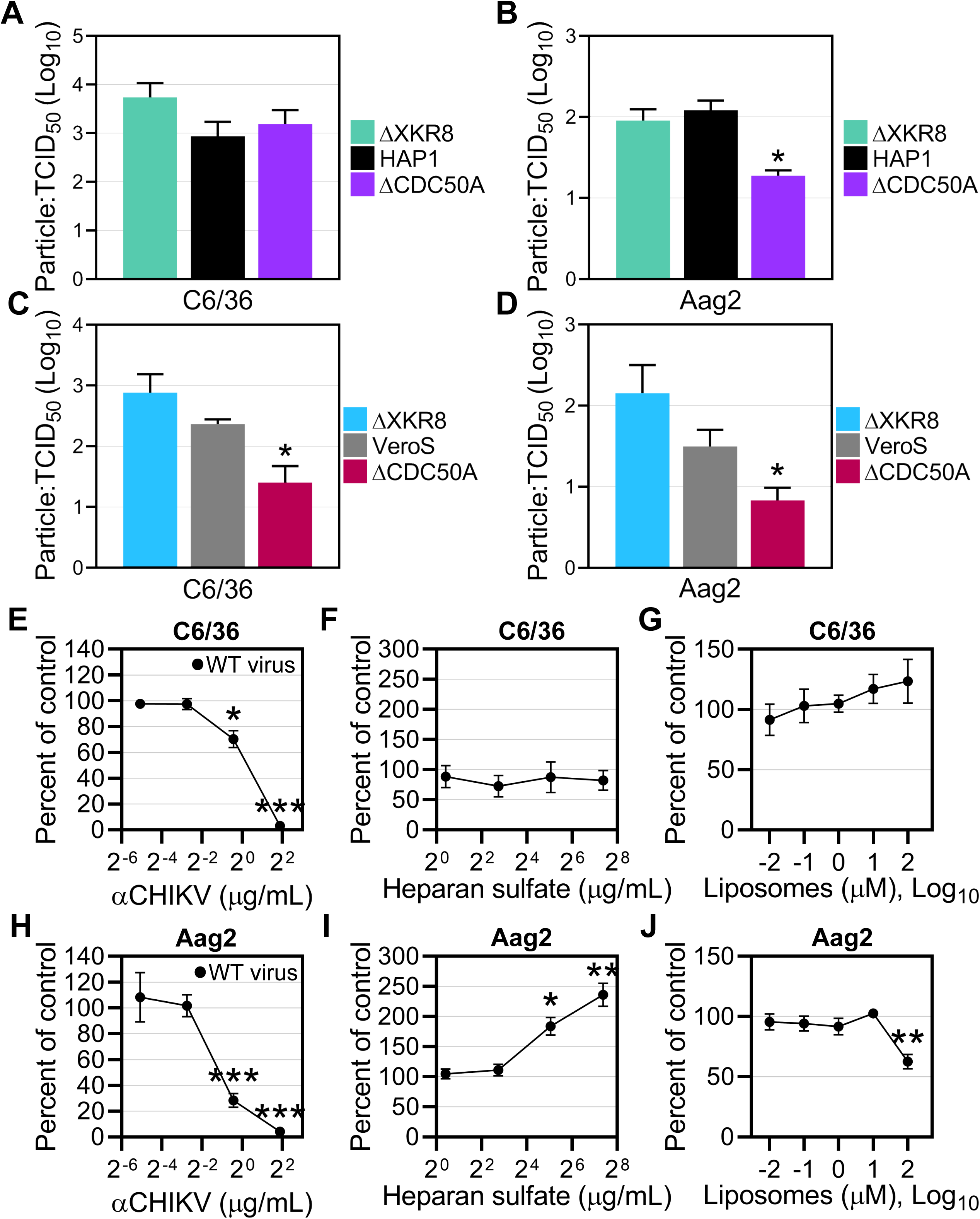
CHIKV viral apoptotic mimicry is cell-type dependent in mosquito cells. The particle (genome copy equivalents) per mL to TCID_50_ units per mL ratio for each sample was used to assess the infectivity of particles produced from (**A, B**) HAP1 cell lines or (**C, D**) VeroS cell lines on mosquito **(A, C**) C6/36 and (**B, D**) Aag2 cells. Congruent with TCID50 methods, equal volumes of collected virus were added to each cell line in the panel without inoculum removal and scored based on CPE and GFP^+^. At least three independent replicates were conducted with bars representing the mean and error (±SEM). Infectivity values were natural log (ln) transformed prior to performing an unpaired parametric student T-test relative to parental cells. (**E-J**) CHIKV-N*luc* stocks were used to assess the ability of (**E, H**) CHIKV antibody, (**F, I**) heparan sulfate, or (**G, J**) PS-containing liposomes (30% PS: 69% PE: 1% PC), to block infections into either C6/36 or Aag2 mosquito cells at the indicated concentrations. Twenty-four hours following infection the cells were lysed with NanoGlo substrate and lysates were quantified with a GloMax Explorer. At least three independent replicates were conducted with dots representing means and error bars representing ±SEM. Values of significance are shown as * (*p* < 0.05), ** (*p* < 0.01), *** (*p* < 0.001).

CHIKV infection in C6/36 cells was inhibited with neutralizing α-CHIKV antibody (**Figure 7E**) as expected, but unaffected by the addition of either heparan sulfate (**Figure 7F**) or liposomes (**Figure 7G**). CHIKV infection in Aag2 cells was also blocked by α-CHIKV antibody (**Figure 7H**) and, in contrast to C6/36 cells, infection was enhanced with heparan sulfate addition (**Figure 7I**). We observed a reduction in CHIKV infection in Aag2 cells treated with our highest concentration (100 µM) of liposomes (**Figure 7J**), congruent with the enhanced infectivity in Aag2 cells with particles produced in flippase KO cells (**Figure 7B, D**). These data support that viral apoptotic mimicry can also enhance CHIKV infection in mosquito cells, but in a cell-type dependent manner.

## Discussion

In this study, we demonstrate that CHIKV entry into mammalian Vero cells and mosquito Aag2 cells is enhanced through viral apoptotic mimicry, a process that involves PS on the virion envelope binding to PSRs on the cellular surface. However, CHIKV infection in mammalian HAP1 and NIH3T3 cells or mosquito C6/36 cells was not affected by virion-associated PS levels, indicating that viral apoptotic mimicry for CHIKV is cell-type dependent (**Figure 8**).

**Figure 8.**
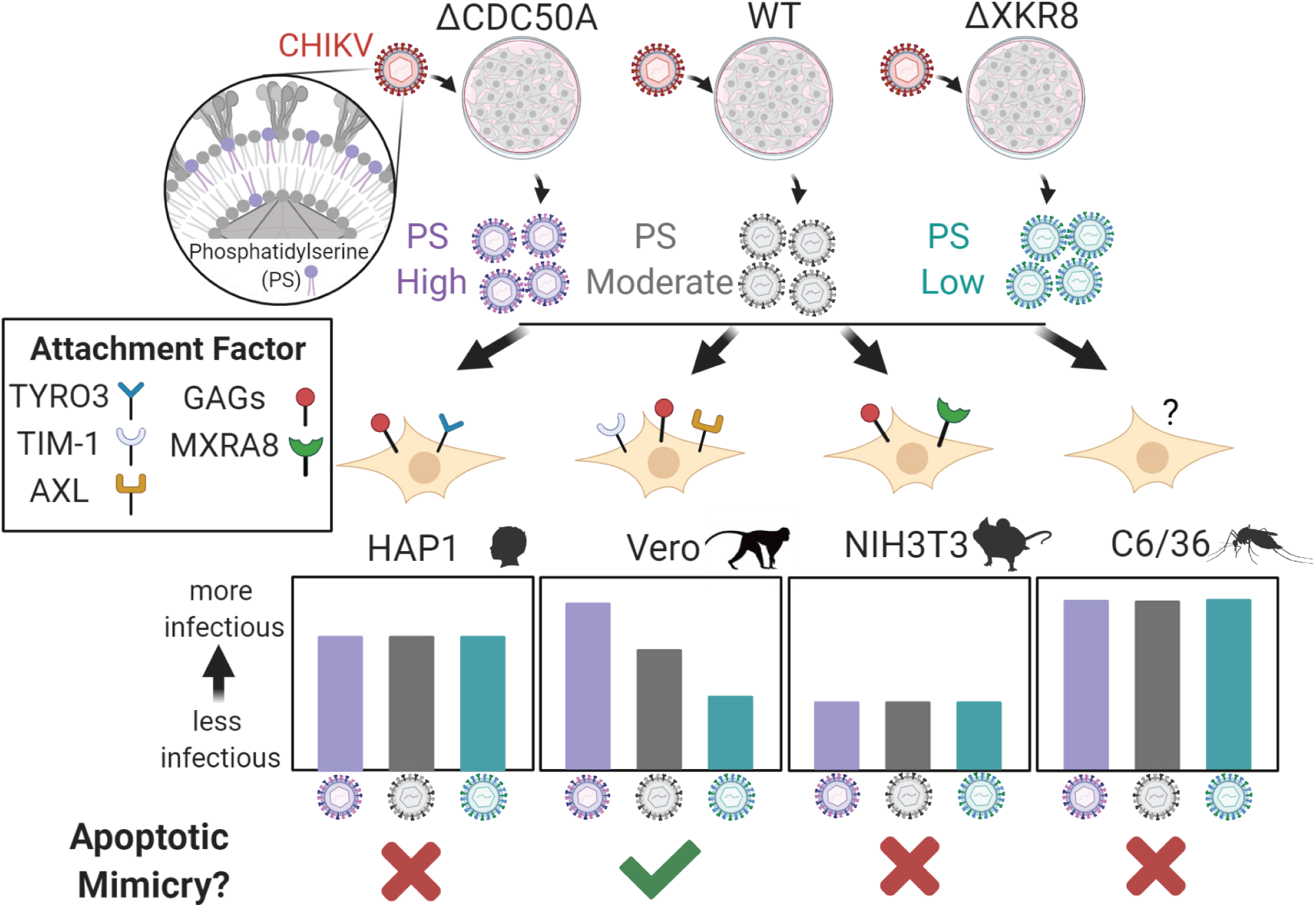
CHIKV viral apoptotic mimicry is cell-type dependent. A visual summary of the current study and main findings. To study the role of viral apoptotic mimicry in CHIKV infection we generated CHIKV virions with distinct levels of phosphatidylserine (PS) exposed on the viral envelope. First, HAP1 and VeroS cells were genetically modified by creating either a flippase subunit KO line (ΔCDC50A) or a scramblase KO line (ΔXKR8) to exploit natural phospholipid translocation dynamics. Abolishing flippase activity (ΔCDC50A) resulted in cells with constitutively high levels of externalized PS in the plasma membrane, whereas impairing scramblase activity (ΔXKR8) resulted in low levels of PS. Next, CHIKV virus was passaged once through each genetically modified cell line to produce viral particles with discrete levels of PS in the outer leaflet of the virion envelope. These produced particles were placed on several cell lines with unique combinations of identified CHIKV attachment factors to assess the role of viral apoptotic mimicry during CHIKV infection in both mammalian (HAP1, Vero, and NIH3T3) and mosquito (C6/36 and Aag2) cells. We found the role of viral apoptotic mimicry in CHIKV infection to occur in a cell type dependent manner. Our data support that GAGs such as heparan sulfate are a major contributor to infection in HAP1 cells, while the PSR TYRO3 is not. CHIKV PS levels correlated with infectivity in Vero cells containing the PSRs TIM-1 and AXL, providing a role of viral apoptotic mimicry in CHIKV entry. Further, we found CHIKV infectivity on NIH3T3 cells to be unaffected by virion PS levels. Lastly, we did not observe a clear role of PS dependent entry in mosquito C6/36 cells, but CHIKV infection in mosquito Aag2 cells was enhanced by viral apoptotic mimicry.

The efficiency of CHIKV entry depends on which attachment factors are present. HAP1 cells endogenously produce GAGs (26) and the PSR TYRO3 (**Figure 5**). Our data supports that CHIKV attachment to HAP1 cells occurs primarily through GAGs, as previously shown (26), and is not PS-dependent. In contrast to HAP1 cells, Vero cells naturally produce the PSRs TIM-1 and AXL (47) and virion-associated PS enhances CHIKV infection in these cells. Further, PS dependence was not observed to enhance CHIKV entry in mouse fibroblast NIH3T3 cells, which display the nonessential proteinaceous receptor MXRA8 (22). The identity of CHIKV attachment factors in the mosquito vector remains unresolved. Neither GAGs nor PS appear involved in CHIKV infection in mosquito C6/36 cells. However, virions containing high amounts of accessible PS produced from CDC50A KO cells were more infectious on Aag2 cells and PS-containing liposomes inhibited CHIKV infection by 38%.

We verified that CHIKV acquires a particle envelope with a lipid bilayer composition reflective of the cellular membrane from which it buds. Virions budding late in infection are enriched in external PS as infected cells become apoptotic. These late produced PS-rich virions can demonstrate improved infectivity through binding to nearby susceptible cells via PSRs. Thus, apoptotic induction during CHIKV infection has pro-viral effects through improved attachment efficiency. Viral apoptotic mimicry likely contributes to the broad host and cellular tropism of CHIKV, but the relevance of this attachment mechanism in viral establishment, dissemination, and transmission between humans and mosquitoes has yet to be determined *in vivo*.

Prior characterization of the regulation of phospholipid distribution within the plasma membrane (PM) lipid bilayer enabled us to alter natural phospholipid dynamics through the deletion of key cellular proteins. These data confirm that knocking out flippase activity results in the accumulation of PS on the outer leaflet of the PM. Further, low levels of PS are maintained on the extracellular side of the lipid bilayer in scramblase KO HAP1ΔXKR8 cells regardless of whether the cell is healthy or apoptotic. Similar amounts of external PS between the parental VeroS cells and our VeroSΔXKR8 scramblase KO line even after CHIKV infection suggest that VeroS cells contain low scramblase activity. A similar study examining Ebola particle infectivity found that XKR8 was primarily distributed in Vero cell cytoplasmic membranes rather than the PM, which may explain reduced PS scrambling on Vero cell surfaces (61).

Considering the prominent role of the PM in CHIKV replication, we investigated whether viral kinetics, as opposed to particle infectivity, was altered in our flippase and scramblase KO cell lines. CHIKV spread through the HAP1 cell lines at similar rates and produced similar levels of viral progeny as evidenced by comparable levels of infectious particles when titrated on HAP1 cells. Preventing PS accumulation on the exterior of the host cell, as in the context of our scramblase KO HAP1ΔXKR8 line, did not appear to affect CHIKV replication kinetics. This suggests that in the absence of PSR attachment, CHIKV can efficiently use alternative attachment factors to bind to HAP1 cells. Indeed, we found the glycosaminoglycan (GAG) heparan sulfate, competitively inhibited CHIKV infection in HAP1 cells, supporting previous findings (26). Interestingly, the boost in CHIKV infection with exogenous ectopic MXRA8 on HAP1 cells suggests that even in the presence of native GAGs, the addition of a proteinaceous receptor aids viral attachment efficiency, while TIM-1 addition did not (**Figure 1C**).

In contrast, CHIKV propagated through CDC50A KO cells produced particles that were more infectious than those derived from WT or ΔXKR8 cells when titrated on VeroS cells. Particles produced in ΔCDC50A cells contain higher levels of PS, suggesting CHIKV entry into Vero cells can be promoted through apoptotic mimicry. Moreover, Vero cells lacking PSRs are less susceptible to CHIKV infection, suggesting that in the absence of PSRs on Vero cells the molecular components facilitating virion attachment and entry are inefficient. Interestingly, the presentation of MXRA8 or additional TIM-1 on VeroS cells did not enhance CHIKV infection, suggesting that the endogenous PSRs efficiently mediate CHIKV entry (**Figure 1D**). These data further support the importance of affinity between virus-cell interactions in CHIKV entry.

NIH3T3 cells were the only cells examined which endogenously present MXRA8 on the cellular surface (22). CHIKV infection in NIH3T3 cells was unaffected by viral envelope PS levels (**Figure 6A, B**). MXRA8 is a nonessential proteinaceous receptor of CHIKV (22, 34, 62) that binds to the E2 glycoprotein (63, 64), whereas TIM-1 binds PS lipids (48) on the virion envelope. MXRA8 may promote stronger CHIKV cellular attachment compared to TIM-1 due to either greater interaction accessibility based on virion structure or a stronger binding affinity. While recent work has established that NIH3T3 cells have a low abundance of MXRA8 on the cellular surface relative to human U2OS cells (26), the overall avidity between MXRA8 and CHIKV-E is likely stronger than those between PSRs and virion-PS; negating any PS-dependent attachment enhancement in NIH3T3 cells.

It is well documented that the attenuated CHIKV strain used in this study (181/c25) displays increased GAG dependence compared to circulating pathogenic strains based on interactions with residue 82 on E2 (25, 26, 65–67). The degree of GAG dependence appears to be strain specific (26). Given that we observed viral apoptotic mimicry in Vero cells using a CHIKV strain with strong GAG affinity, suggests that endemic strains may either (i) be more reliant on alternative attachment factors such as PSRs and/or (ii) be less infectious in the same context. A recent study found increased CHIKV infectivity with East-Central-South-African (ECSA) strain LR2006-OPY1 and West African (WA) strain 37997 relative to Asian strain 181/c25 in 293T cells stably expressing TIM-1 or a TIM-1 variant lacking the cytoplasmic domain (32).

In humans, CHIKV infection is initiated by virion deposition into the skin dermis during the bite of an infectious female mosquito. Fibroblasts, keratinocytes, and resident macrophages support initial CHIKV infection (22, 68). While fibroblasts are permissive for CHIKV, the infection appears to be predominately MXRA8-dependent (22). Keratinocytes present in the basal layer of the skin epidermis express both TIM-1 and AXL (50, 69) and are susceptible to CHIKV infection (22). Interestingly, an immortalized keratinocyte cell line (HaCat) was shown to be refractory to CHIKV infection due to an induction of interferon (68). However, a more recent study demonstrated that HaCat cells produced low levels of AXL along with undetectable levels of TIM-1, and that the addition of TIM-1 increased CHIKV susceptibility and permissivity (32). Thus, keratinocytes may have a larger role in CHIKV infection establishment *in vivo* than previously thought. Macrophages also display PSRs, conferring phagocytic properties of apoptotic body clearance (70–72). PS-rich virions from either infected fibroblasts, keratinocytes, or mosquito inoculation may serve as an ideal target to attach to PSRs on resident macrophages. While macrophage infection via apoptotic mimicry could facilitate CHIKV dissemination *in vivo*, macrophages often are poor producers of CHIKV virus *in vitro* (73).

Current evidence suggests that the long-term arthralgia associated with CHIKV infection is due to immune-mediated tissue pathology (74, 75). While the cellular surface abundance of MXRA8 overlaps with tissue types of pathogenic relevance (22, 34, 64, 74) and MXRA8 deficient mice have reduced joint pathology (34), apoptotic mimicry may promote initial infection or dissemination as CHIKV infection *in vivo* still proceeds in MXRA8-deficient mice (34). Future research should aim to understand the role of apoptotic mimicry in the context of infection establishment and dissemination using *in vivo* model systems.

The transmission of CHIKV to a mosquito vector can occur when a susceptible mosquito ingests blood from a viremic mammalian host. Key differences exist between mammalian-made and mosquito-made virions that affect infectivity (76–82). For example, differences in protein post-translation modifications between mammalian and invertebrate cells contribute to differences in virion infectivity (N-glycosylation (80, 83)). In addition, the plasma membrane of insect cells has a distinct lipid profile from that of mammalian cells (60, 84). While we did not see strong support for viral apoptotic mimicry influencing CHIKV infection in mosquito C6/36 cells, CHIKV demonstrated PS-dependent enhancement in mosquito Aag2 cells. In Aag2 cells, we observed that elevated externalized PS levels increased CHIKV particle infectivity and PS-liposomes inhibited CHIKV infection. Given the clear role of cell type in dictating PS dependence, it may not be appropriate to conclude the role of viral apoptotic mimicry in mosquitoes from either C6/36 cells or Aag2 cells, which are not representative of the cell types implicated in natural infection. Cell factors that facilitate CHIKV entry into mosquito cells remain elusive, however, mosquitoes do not produce homologs to MXRA8 (34), drosophila encode PSR orthologs, and apoptotic cell clearance via phosphatidylserine exposure is conserved (85).

The examination of CHIKV entry efficiency throughout infection *in vivo* is needed to delineate the relevance of attachment via PSRs among other surface molecules including GAGs and MXRA8 in influencing infection establishment, dissemination, and cross-species transmission. Future studies should aim to robustly characterize the role of each CHIKV attachment factor, including viral apoptotic mimicry, in systems that more closely resemble natural infection.

## Materials & Methods

### Cell lines

Human near-haploid cells (HAP1) derived from the male chronic myelogenous leukemia cell line KBM-7, HAP1 flippase KO line (HAP1ΔCDC50A, HZGHC005423c007), and HAP1 scramblase KO line (HAP1ΔXKR8, HZGHC005916c007) were purchased from Horizon Discovery (United Kingdom). HAP1 and HAP1 KO lines were cultured in Iscove’s modified Dulbecco’s medium (IMDM) supplemented with 8% (v/v) fetal bovine serum (FBS). All vervet monkey cells (VeroS, VeroSΔCDC50A, VeroSΔXKR8, Vero, and VeroΔPSR,) were maintained with DMEM supplemented with 5% (v/v) FBS. The Vero and VeroΔPSR cells were a kind gift from Dr. Wendy Maury at the University of Iowa (47). Mouse embryo fibroblasts cells (NIH3T3) were purchased from ATCC (CRL-1658) and maintained with DMEM supplemented with 10% (v/v) FBS. All mammalian cells were kept in a humidified chamber held at 37°C and with a 5% CO_2_ content. Mosquito *Aedes albopictus* C6/36 (ATCC, CRL-1660) were maintained with Leibovitz’s L-15 medium supplemented with 10% (v/v) FBS in a humidified chamber held at 28°C. *Aedes aegypti* Aag2 (ATCC, CCL-125) larval homogenate cells were maintained in SFX insect medium with 2% (v/v) FBS in a humidified chamber at 28°C.

### CRISPR-Cas9 mediated generation of VeroS KO cell lines

Three guide RNAs targeting each *Chlorocebus sabaeus* gene, XKR8 (GGCACTGCTCGACTACCACC, TGATCTACTTCCTGTGGAAC, CAGCTATGTGGCCCTGCACT) and CDC50A (TACGGCTGGCACGGTGCTAC, TCGTCGTTACGTGAAATCTC, GTGAACTGGCTTAAACCAGT), were inserted into pSpCas9(BB)-2A-GFP (pX458), which was a gift from Feng Zhang (Addgene plasmid #48138) (86) and verified using Sanger sequencing. VeroS cells were transfected with equivalent amounts of pSpCas9(BB)-2A-GFP bearing each of the three guide RNAs using GeneJuice (Sigma-Aldrich, cat. 70967). Three days post-transfection, VeroS cells were counted and distributed at a density of 0.5 cells per well into 96-well plates. Cells were monitored for 3 weeks to maintain single colony clones, and non-clonal wells were discarded. Wells corresponding to single clones were expanded to 24-well plates and assessed for CRISPR knockout. CRISPR XKR8 and CDC50A KOs were validated by extracting total DNA and PCR amplifying the guide RNA targeted regions. PCR amplicons spanning *xkr8* CRISPR regions were gel purified and submitted for Sanger sequencing to verify *xkr8* modification, which showed a 136 bp deletion in exon 2. We could not amplify *cdc50a* CRISPR regions in exons 1 and 3 but could amplify the CRISPR region targeting exon 5, indicating the presence of a large deletion spanning multiple exons in CDC50A. CRISPR CDC50A KO was also validated using a functional screen for externalized PS. Cells were washed in PBS and the media was changed to Annexin V binding buffer (1mM HEPES pH 7.4, 14 mM NaCl, 0.25mM CaCl_2_) with Annexin V-PE conjugate at 1:50 (v/v) and incubated at room temperature for 15 mins. Cells were washed with Annexin V binding buffer and visually analyzed under a fluorescence microscope. Cells with high PS staining compared with parental cells were expanded for future experiments. Parental cells were treated with 1 mM MG132 for 2 hrs were used as a positive signal control.

### DNA transfections

Transfection efficiency and cytotoxicity varied with each cell line and gene KO. We paired different transfection reagents with different cell lines to optimize transfection efficiency and reduce cytotoxicity. All HAP1 cells were transfected with JetOptimus (PolyPlus, cat. 117-07), VeroS, VeroSΔXKR8 and 293T cells with GeneJuice (Sigma-Alrich, cat. 70967), and CDC50A KO lines with Viafect (Promega, cat. E4981) according to manufacturer recommendations. Expression vectors encoding a GFP-fused transmembrane hTIM-1 (a gift from Wendy Maury at the University of Iowa), pCS6-L-SIGN (Transomic; cat. BC038851), pTiger-MXRA8, or pCMV-GFP were used to assess CHIKV virion surface binding kinetics.

### Viruses

Chikungunya virus (CHIKV) strain 181 clone 25 (181/c25) was used to conduct experiments in a BSL2 laboratory environment. Full-length DNA CHIKV clones containing reporter genes (*gfp, mKate, or Nluc*) were linearized and *in vitro* transcribed (Ambion, cat. AM1344) adhering to the manufacturer’s protocol. Infectious CHIKV virions expressing reporter genes were recovered after direct RNA transfection (1µg) into VeroS cells with Lipofectamine 3000 (Thermofisher, cat. L3000001). Unless otherwise stated, viral stocks were propagated in VeroS cells and passage 3 viral stocks were used for all experiments. The amount of infectious virus was determined by calculating the 50% tissue culture infective dose (TCID_50_) units per mL through end-point dilution using the Spearman-Karber method (87). Replication deficient adenovirus (Ad5CMVeGFP) was purchased from UI Viral Vector Core Web at a predetermined high titer of 5×10^10^ PFU/mL.

### Real-time quantification PCR (RT-qPCR) of genome equivalents

CHIKV genome equivalents/mL were calculated via RT-qPCR. Viral RNA was extracted from infected cell supernatant (Zymo, cat. 11-355), eluted in nuclease-free water, and converted to cDNA with random hexamers (ThermoFisher, cat. 4388950) following kit protocols. RT-qPCR reactions were set up with cDNA, TaqMan Gene Master Mix (Applied Biosystems, cat. 4369016), primers, and TaqMan probe (5’-6FAMACTTGCTTTGATCGCCTTGGTGAGAMGBNFQ-3’) as previously described (88) with each sample run in duplicate. A plasmid-based standard curve of a full-length CHIKV clone was used to enumerate the total number of genome equivalents per mL of the original sample. A no template control (NTC) and no amplification control (NAC) were included in each run on a StepOne platform (Applied Biosystems, cat. 4376357). The amplification profile included 1 cycle of 2 mins at 50°C, 10 mins at 95°C, followed by 40 cycles of 15 secs at 95°C and 1min at 60°C.

### 293T immunofluorescence staining

293T cells were plated at 2.0×10^5^ cells per well in 12-well plates 48 hrs before immunofluorescence staining. Cells were transfected with plasmids encoding MXRA8, hTIM-1-GFP, or L-SIGN along with a plasmid encoding GFP 24 hrs before immunofluorescence staining. Transfected cells were rapidly cooled and stained in blocking solution (dPBS with 2% (v/v) bovine serum albumin (BSA)) containing anti-MXRA8 (1:100, W040-3, MBL International), anti-hTIM1(1:100, AF1750, R&D Systems), or anti-CLEC4M (L-SIGN/CD299) 2G1 antibody (1:100, MA5-21012, Thermo) at 4°C with gentle shaking for 1 hr. Cells were washed with PBS before lifting the cells with a scraper. Cells were pelleted (500xg for 5 mins), resuspended, and washed in PBS two additional times before adding secondary anti-goat Cy5 (1:2500, 072-02-13-06, KPL) or anti-mouse Alexa Fluor 647 (1:2500, A32728, Invitrogen) and incubated at 4°C in the dark for 30 mins. Cells were washed with PBS three times and then analyzed via flow cytometry. Cell populations were gated using forward scatter/side scatter. The mean fluorescence intensity (MFI) of a minimum of 10000 GFP positive cells were quantified per experimental condition and performed three independent times. Secondary only and GFP only transfected cells were stained with primary and secondary antibodies and used as controls for all conditions. All cells were analyzed using a NovoCyte Quanteon (Aligent) flow cytometer.

### Quantification of cellular outer leaflet phosphatidylserine (PS)

Cellular surface levels of PS were assessed using Promega’s RealTime-Glo Annexin V Apoptosis and Necrosis Assay (Promega, cat. JA1012) according to manufacturer specifications. HAP1 or VeroS cell lines were plated in media supplemented with 0.1 M HEPES at 3.0×10^4^ or 10^4^ cells per well, respectively, in a 96-well black-walled, clear bottom plate 1 day prior to treatment. Cells were infected with CHIKV-*mKate* (MOI of 1.0) or mock infected. Kit components 1-4 were added to cells 1 hr following infection and the plate was moved into a pre-warmed GloMax Explorer. Kit components 1-4 were used when assaying HAP1 cells at 0.5x concentration as cytotoxicity was observed at 1x manufacturer recommendations. Automated luminescence (Annexin V) and fluorescence (membrane integrity) measurements were collected every 30 mins in a GloMax Explorer (Promega) held at 37°C.

### Quantification of viral outer leaflet phosphatidylserine (PS)

#### Virus Production

T75 flasks were seeded with wild type, ΔXKR8, and ΔCDC50A HAP1 and VeroS cells with 7.2 x 10^6^ cells or 3.6 × 10^6^ cells, respectively. After 24 hrs, wild-type and ΔXKR8 cells were infected with CHIKV using MOI =0.001 and ΔCDC50A cells were infected using MOI=0.01. After 12 hrs at 37°C, inoculum was removed, cells were treated with citric acid buffer (40 mM citric acid, 10 mM KCl, 135 mM NaCl [pH 3.0]) for 1 min, rinsed, and FBS-free media was added. After incubating for an additional 36 hrs, the supernatant was collected, cleared twice using centrifugation (6,000xg) and overlaid on a 20% sucrose cushion. Overlaid supernatants were then subjected to ultracentrifugation at (234,116xg) for 2 hrs at 4°C. Pellets were resuspended in 100 μL PBS.

#### Input normalization

Prior to staining, purified CHIKV samples were normalized using RT-qPCR: To detect CHIKV capsid (C) levels, normalized samples were denatured using SDS-urea buffer (200 mM Tris [pH 6.8], 8 M urea, 5% SDS, 0.1 mM EDTA, 0.03% bromophenol blue), run on Mini-PROTEAN TGX Stain-Free Precast Gels (Bio-Rad), and imaged with a ChemiDoc XRS digital imaging system (Bio-Rad). Gels were then subjected to immunoblot analysis for CHIKV E using an anti-E antibody (1:1000, R&D Systems, MAB97792SP).

#### Particle surface PS staining

Equivalent numbers of CHIKV particles were conjugated to 4-μm aldehyde/sulfate latex beads (Thermo Fisher Scientific) overnight at 4°C with gentle shaking. Due to differences in viral yields between cell types, beads were bound with approximately 10^6^ genome equivalents from HAP1 cell lines and 10^9^ genome equivalents from VeroS cell lines. Beads were blocked with a final concentration of 1% (v/v) bovine serum albumin (BSA) in PBS for 2 hrs while rotating at room temperature. Beads were washed 3 times with 1% (v/v) BSA in PBS and then incubated with 100 μl of AnV binding buffer containing AnV-PE conjugate for 30 mins on ice. Beads were diluted 1:4 in AnV binding buffer and analyzed using the NovoCyte Quanteon flow cytometer (Aligent). Bead only samples were included as a mock control.

### Multi-step CHIKV replication kinetics

A low MOI (0.01) was used to assess replication kinetics over multiple replication cycles. Cells were seeded at a density of 3×10^5^ cells/mL for HAP1 lines and 2.5×10^5^ cells/mL for VeroS lines and inoculated with virus in FBS-free media for 1 hr at 37°C before removing viral inoculum and replacing with complete media. At each time point, the supernatant was collected, and fresh media was replaced. Supernatants were frozen at -80°C until all time-points were collected. Virus was titrated on the indicated cell line and the number of infectious particles per mL was calculated by determining the 50% tissue culture infective dose (TCID_50_) as described above.

### Cell-to-cell viral spread kinetics

Cells were plated at either 7.5×10^4^ cells per well in a 48-well plate (HAP1 lines) or 5×10^4^ cells per well in a 24-well plate (Vero lines) 1 day prior to infection. Assuming cells doubled overnight, cells were inoculated with CHIKV-GFP virus (MOI of 0.1). After 1 hr (T = 0 hpi) virus inoculum was removed and replaced with complete media. At the indicated time, cells were lifted in trypsin, resuspended in PBS, and fixed in 1.5% (v/v) formaldehyde. GFP positive cells were enumerated in a NovoCyte Quanteon (Aligent) flow cytometer. Cells were first gated based on forward/side scatter, and cellular aggregates were removed by gating with forward scatter area to height. Uninfected cells were used to set the GFP gate. 10000 live cells were collected and the percent infection (% GFP+) was recorded and compared over time.

### Entry assays

Viruses expressing *gfp* were used to determine if cellular attachment and internalization varied across cell lines. 7.5×10^4^ cells per well were plated in a 48-well format for HAP1 cell lines and 5×10^4^ cells per well in a 24-well format for Vero cell lines 1 day prior to infection. A high MOI was used to infect ∼50% of the cell population in the initial round of infection. Virus inoculum was added to cells for 1 hr at 37°C after which viral inoculum was removed and replaced with complete medium, marking 0 hpi. CHIKV requires low pH in the endosome for membrane fusion and genome release. To ensure we were measuring GFP production from only the first round of infection for CHIKV, 30 mM ammonium chloride (NH_4_Cl) was added 2 hpi or with virus inoculum to demonstrate blocking potency. Cells were lifted at either 12 hpi (CHIKV) or 20 hpi (AdenoV), resuspended in PBS, and fixed with 4% formaldehyde. A NovoCyte Quanteon (Aligent) flow cytometer was used to determine the percentage of GFP^+^ cells. Cell gating was the same as described above for viral spread.

### Surface biotinylation

Parental HAP1 and KO lines were seeded at 5×10^5^ cells per well in a 6-well plate and incubated for 48 hrs. Cells were then washed with cold PBS and biotinylated with 0.5 mg/ml sulfosuccinimidyl-2-(biotinamido) ethyl-1,3-dithiopropionate (ThermoFisher) while gently shaking for 45 mins on ice. The reaction was quenched using Tris-HCl. Cells were lysed in M2 lysis buffer (50 mM Tris [pH 7.4], 150 mM NaCl, 1 mM EDTA, 1% Triton X-100) at 4°C and clarified with centrifugation at 17,000Xg for10 min. To purify surface exposed proteins, lysates were incubated with streptavidin sepharose beads (GE Healthcare) overnight while rotating at 4°C. Following incubation, the streptavidin sepharose beads were washed in buffer 1 (100 mM Tris, 500 mM lithium chloride, 0.1% Triton X-100) and then in buffer 2 (20 mM HEPES [pH 7.2], 2 mM EGTA, 10 mM magnesium chloride, 0.1% Triton X-100), incubated in urea buffer (200 mM Tris [pH 6.8], 8 M urea, 5% sodium dodecyl sulfate [SDS], 0.1 mM EDTA, 0.03% bromophenol blue, 1.5% dithiothreitol [DTT]) for 30 min at 56°C, and subjected to immunoblot analysis using anti-βactin C4 (1:1000, Santa Cruz Biotechnology, sc-47778) and anti-Tyro-3 (1:1000, R&D Systems, MAB859100) antibodies. Immunoblots were probed with appropriate secondary antibodies conjugated with HRP and imaged with a ChemiDoc XRS digital imaging system (Bio-Rad).

### Vero immunofluorescence staining

Vero cell lines were plated at 10^5^ cells per well in a 24-well format 24 hrs before immunofluorescence staining. Cells were rapidly cooled by placing cells on ice and replacing media with ice cold PBS for 15 mins. PBS was removed and replaced with 200 µL of blocking solution (dPBS with 2% (v/v) BSA) containing α-TIM1(1:100, AF1750, R&D Systems) or α - AXL antibody (1:100, AF154, R&D Systems) and incubated at 4°C and gently shook for 1 hr. Cells were washed three times with PBS. Following the third wash, 200 µL of blocking solution containing α -goat Cy5 (1:2500, 072-02-13-06, KPL) and incubated at 4°C in the dark and gently rocked for 30 mins. After three PBS washes, cells were scraped, homogenized, and analyzed via flow cytometry. Cell populations were gated using forward scatter/side scatter. The mean fluorescence intensity (MFI) of a minimum of 5,000 live cells was quantified per experimental condition. Secondary only was used as a negative control in all conditions. All cells were analyzed using a NovoCyte Quanteon (Aligent) flow cytometer.

### Particle infectivity

We used the ratio of genome equivalents to infectious viral particles to assess particle infectivity. This ratio represents the number of particles needed to start an infection. A smaller value indicates a virus stock is more infectious, or each particle has a higher probability of starting an infection. Particle number was determined by quantifying the number of genome equivalents in the virus preparation using qRT-PCR described above. Infectivity was determined by TCID_50_ units per mL. CHIKV readily forms plaques on VeroS cells, however, not all our cell lines tolerated forming a confluent monolayer under an agar overlay and therefore TCID_50_ units were used when comparing various cell lines.

### Competition assays

CHIKV-N*luc* stocks were used to assess the ability of antibody, liposomes, or heparin sulfate to block infections into the indicated cell lines. HAP1 cells were seeded at 5×10^4^ cells per well in a 96-well plate 1 day prior to infection. For each well in the competition assay approximately 150 CHIKV-N*luc* virions were added. 24 hrs following infection, cells were lysed with NanoGlo substrate and lysates were quantified in a GloMax Explorer (Promega) according to manufacturer’s instructions for all competitive inhibition assays.

*Antibody competition*: virus was incubated with the indicated concentrations of CHIKV polyclonal antibody (IBT, cat. 04-008) or no antibody control at room temperature for 45 mins. After incubation, the virus-CHIKV antibody mix was added to cells.

*Heparan competition*: the indicated concentration of heparan sulfate (Sigma-Aldrich, cat. H7640-1MG), or control PBS was added to cells at 37°C for 10 mins prior to infection. After the 10 min pre-treatment, virus was added.

*Liposome competition*: the indicated concentration of liposomes (30% PS: 69% PE: 1% PC) (48) or PBS was added to cells at 37°C for 10 mins prior to infection. After the 10 min pre-treatment, virus was added.

### Statistical analysis

Data were visualized and analyzed using GraphPad Prism software (v8.2.1, windows 64-bit). An unpaired parametric Student T-test assuming equal variance was used to test for statistical significance for data on a linear scale (e.g., percent infected). An unpaired parametric Student T-test using a Welch’s correction was used to test for statistical significance for normalized data (e.g., relative infection, normalized MFI). Logarithmic data were natural log (ln) transformed and then assessed with an unpaired parametric Student T-test assuming equal variance (e.g., genome equivalents, specific infectivity).

## Acknowledgements

We would like to thank James Barber and the CVM flow cytometer core at the University of Georgia for their technical assistance.

## References

1. Gerardin P, Guernier V, Perrau J, Fianu A, Le Roux K, Grivard P, Michault A, de Lamballerie X, Flahault A, Favier F. 2008. Estimating Chikungunya prevalence in La Reunion Island outbreak by serosurveys: two methods for two critical times of the epidemic. BMC Infect Dis 8:99.

2. Moro ML, Gagliotti C, Silvi G, Angelini R, Sambri V, Rezza G, Massimiliani E, Mattivi A, Grilli E, Finarelli AC, Spataro N, Pierro AM, Seyler T, Macini P, Chikungunya Study G. 2010. Chikungunya virus in North-Eastern Italy: a seroprevalence survey. Am J Trop Med Hyg 82:508–11.

3. Brighton SW, Prozesky OW, de la Harpe AL. 1983. Chikungunya virus infection. A retrospective study of 107 cases. S Afr Med J 63:313–5.

4. Sissoko D, Malvy D, Ezzedine K, Renault P, Moscetti F, Ledrans M, Pierre V. 2009. Post-epidemic Chikungunya disease on Reunion Island: course of rheumatic manifestations and associated factors over a 15-month period. PLoS Negl Trop Dis 3:e389.

5. Economopoulou A, Dominguez M, Helynck B, Sissoko D, Wichmann O, Quenel P, Germonneau P, Quatresous I. 2009. Atypical Chikungunya virus infections: clinical manifestations, mortality and risk factors for severe disease during the 2005-2006 outbreak on Reunion. Epidemiol Infect 137:534–41.

6. Tsetsarkin KA, Vanlandingham DL, McGee CE, Higgs S. 2007. A single mutation in chikungunya virus affects vector specificity and epidemic potential. PLoS Pathog 3:e201.

7. Agarwal A, Sharma AK, Sukumaran D, Parida M, Dash PK. 2016. Two novel epistatic mutations (E1:K211E and E2:V264A) in structural proteins of Chikungunya virus enhance fitness in Aedes aegypti. Virology 497:59–68.

8. Powers AM, Brault AC, Tesh RB, Weaver SC. 2000. Re-emergence of Chikungunya and O’nyong-nyong viruses: evidence for distinct geographical lineages and distant evolutionary relationships. J Gen Virol 81:471–9.

9. Volk SM, Chen R, Tsetsarkin KA, Adams AP, Garcia TI, Sall AA, Nasar F, Schuh AJ, Holmes EC, Higgs S, Maharaj PD, Brault AC, Weaver SC. 2010. Genome-scale phylogenetic analyses of chikungunya virus reveal independent emergences of recent epidemics and various evolutionary rates. J Virol 84:6497–504.

10. Angelini R, Finarelli AC, Angelini P, Po C, Petropulacos K, Silvi G, Macini P, Fortuna C, Venturi G, Magurano F, Fiorentini C, Marchi A, Benedetti E, Bucci P, Boros S, Romi R, Majori G, Ciufolini MG, Nicoletti L, Rezza G, Cassone A. 2007. Chikungunya in north-eastern Italy: a summing up of the outbreak. Euro Surveill 12:E071122 2.

11. Rezza G, Nicoletti L, Angelini R, Romi R, Finarelli AC, Panning M, Cordioli P, Fortuna C, Boros S, Magurano F, Silvi G, Angelini P, Dottori M, Ciufolini MG, Majori GC, Cassone A, group Cs. 2007. Infection with chikungunya virus in Italy: an outbreak in a temperate region. Lancet 370:1840–6.

12. Vega-Rua A, Zouache K, Caro V, Diancourt L, Delaunay P, Grandadam M, Failloux AB. 2013. High efficiency of temperate Aedes albopictus to transmit chikungunya and dengue viruses in the Southeast of France. PLoS One 8:e59716.

13. Ryan SJ, Carlson CJ, Mordecai EA, Johnson LR. 2019. Global expansion and redistribution of Aedes-borne virus transmission risk with climate change. PLoS Negl Trop Dis 13:e0007213.

14. Romi R, Severini F, Toma L. 2006. Cold acclimation and overwintering of female Aedes albopictus in Roma. J Am Mosq Control Assoc 22:149–51.

15. Fischer D, Thomas SM, Suk JE, Sudre B, Hess A, Tjaden NB, Beierkuhnlein C, Semenza JC. 2013. Climate change effects on Chikungunya transmission in Europe: geospatial analysis of vector’s climatic suitability and virus’ temperature requirements. Int J Health Geogr 12:51.

16. Jose J, Snyder JE, Kuhn RJ. 2009. A structural and functional perspective of alphavirus replication and assembly. Future Microbiol 4:837–56.

17. Simizu B, Yamamoto K, Hashimoto K, Ogata T. 1984. Structural proteins of Chikungunya virus. J Virol 51:254–8.

18. Sun S, Xiang Y, Akahata W, Holdaway H, Pal P, Zhang X, Diamond MS, Nabel GJ, Rossmann MG. 2013. Structural analyses at pseudo atomic resolution of Chikungunya virus and antibodies show mechanisms of neutralization. Elife 2:e00435.

19. Voss JE, Vaney MC, Duquerroy S, Vonrhein C, Girard-Blanc C, Crublet E, Thompson A, Bricogne G, Rey FA. 2010. Glycoprotein organization of Chikungunya virus particles revealed by X-ray crystallography. Nature 468:709–12.

20. Kielian M. 1995. Membrane fusion and the alphavirus life cycle. Adv Virus Res 45:113–51.

21. Lu YE, Kielian M. 2000. Semliki forest virus budding: assay, mechanisms, and cholesterol requirement. J Virol 74:7708–19.

22. Zhang R, Kim AS, Fox JM, Nair S, Basore K, Klimstra WB, Rimkunas R, Fong RH, Lin H, Poddar S, Crowe JE, Jr., Doranz BJ, Fremont DH, Diamond MS. 2018. Mxra8 is a receptor for multiple arthritogenic alphaviruses. Nature 557:570–574.

23. Tanaka A, Tumkosit U, Nakamura S, Motooka D, Kishishita N, Priengprom T, Sa-Ngasang A, Kinoshita T, Takeda N, Maeda Y. 2017. Genome-Wide Screening Uncovers the Significance of N-Sulfation of Heparan Sulfate as a Host Cell Factor for Chikungunya Virus Infection. J Virol 91.

24. Weber C, Berberich E, von Rhein C, Henss L, Hildt E, Schnierle BS. 2017. Identification of Functional Determinants in the Chikungunya Virus E2 Protein. PLoS Negl Trop Dis 11:e0005318.

25. Sahoo B, Chowdary TK. 2019. Conformational changes in Chikungunya virus E2 protein upon heparan sulfate receptor binding explain mechanism of E2-E1 dissociation during viral entry. Biosci Rep 39.

26. McAllister N, Liu Y, Silva LM, Lentscher AJ, Chai W, Wu N, Griswold KA, Raghunathan K, Vang L, Alexander J, Warfield KL, Diamond MS, Feizi T, Silva LA, Dermody TS. 2020. Chikungunya Virus Strains from Each Genetic Clade Bind Sulfated Glycosaminoglycans as Attachment Factors. J Virol 94.

27. Prado Acosta M, Geoghegan EM, Lepenies B, Ruzal S, Kielian M, Martinez MG. 2019. Surface (S) Layer Proteins of Lactobacillus acidophilus Block Virus Infection via DC-SIGN Interaction. Front Microbiol 10:810.

28. Bucardo F, Reyes Y, Morales M, Briceno R, Gonzalez F, Lundkvist A, Svensson L, Nordgren J. 2020. Genetic polymorphisms in DC-SIGN, TLR3 and TNFa genes and the Lewis-negative phenotype are associated with Chikungunya infection and disease in Nicaragua. J Infect Dis doi:10.1093/infdis/jiaa364.

29. Wintachai P, Wikan N, Kuadkitkan A, Jaimipuk T, Ubol S, Pulmanausahakul R, Auewarakul P, Kasinrerk W, Weng WY, Panyasrivanit M, Paemanee A, Kittisenachai S, Roytrakul S, Smith DR. 2012. Identification of prohibitin as a Chikungunya virus receptor protein. J Med Virol 84:1757–70.

30. Meertens L, Carnec X, Lecoin MP, Ramdasi R, Guivel-Benhassine F, Lew E, Lemke G, Schwartz O, Amara A. 2012. The TIM and TAM families of phosphatidylserine receptors mediate dengue virus entry. Cell Host Microbe 12:544–57.

31. Moller-Tank S, Kondratowicz AS, Davey RA, Rennert PD, Maury W. 2013. Role of the phosphatidylserine receptor TIM-1 in enveloped-virus entry. J Virol 87:8327–41.

32. Kirui J, Abidine Y, Lenman A, Islam K, Gwon YD, Lasswitz L, Evander M, Bally M, Gerold G. 2021. The Phosphatidylserine Receptor TIM-1 Enhances Authentic Chikungunya Virus Cell Entry. Cells 10.

33. Carnec X, Meertens L, Dejarnac O, Perera-Lecoin M, Hafirassou ML, Kitaura J, Ramdasi R, Schwartz O, Amara A. 2016. The Phosphatidylserine and Phosphatidylethanolamine Receptor CD300a Binds Dengue Virus and Enhances Infection. J Virol 90:92–102.

34. Zhang R, Earnest JT, Kim AS, Winkler ES, Desai P, Adams LJ, Hu G, Bullock C, Gold B, Cherry S, Diamond MS. 2019. Expression of the Mxra8 Receptor Promotes Alphavirus Infection and Pathogenesis in Mice and Drosophila. Cell Rep 28:2647–2658 e5.

35. Kay JG, Fairn GD. 2019. Distribution, dynamics and functional roles of phosphatidylserine within the cell. Cell Commun Signal 17:126.

36. van Meer G, Voelker DR, Feigenson GW. 2008. Membrane lipids: where they are and how they behave. Nat Rev Mol Cell Biol 9:112–24.

37. Takatsu H, Tanaka G, Segawa K, Suzuki J, Nagata S, Nakayama K, Shin HW. 2014. Phospholipid flippase activities and substrate specificities of human type IV P-type ATPases localized to the plasma membrane. J Biol Chem 289:33543–56.

38. Papadopulos A, Vehring S, Lopez-Montero I, Kutschenko L, Stockl M, Devaux PF, Kozlov M, Pomorski T, Herrmann A. 2007. Flippase activity detected with unlabeled lipids by shape changes of giant unilamellar vesicles. J Biol Chem 282:15559–68.

39. Andersen JP, Vestergaard AL, Mikkelsen SA, Mogensen LS, Chalat M, Molday RS. 2016. P4-ATPases as Phospholipid Flippases-Structure, Function, and Enigmas. Front Physiol 7:275.

40. Huang W, Liao G, Baker GM, Wang Y, Lau R, Paderu P, Perlin DS, Xue C. 2016. Lipid Flippase Subunit Cdc50 Mediates Drug Resistance and Virulence in Cryptococcus neoformans. mBio 7.

41. Yang F, Huang Y, Chen X, Liu L, Liao D, Zhang H, Huang G, Liu W, Zhu X, Wang W, Lobo CA, Yazdanbakhsh K, An X, Ju Z. 2019. Deletion of a flippase subunit Tmem30a in hematopoietic cells impairs mouse fetal liver erythropoiesis. Haematologica 104:1984–1994.

42. Segawa K, Kurata S, Yanagihashi Y, Brummelkamp TR, Matsuda F, Nagata S. 2014. Caspase-mediated cleavage of phospholipid flippase for apoptotic phosphatidylserine exposure. Science 344:1164–8.

43. Suzuki J, Imanishi E, Nagata S. 2016. Xkr8 phospholipid scrambling complex in apoptotic phosphatidylserine exposure. Proc Natl Acad Sci U S A 113:9509–14.

44. Bricogne C, Fine M, Pereira PM, Sung J, Tijani M, Wang Y, Henriques R, Collins MK, Hilgemann DW. 2019. TMEM16F activation by Ca(2+) triggers plasma membrane expansion and directs PD-1 trafficking. Sci Rep 9:619.

45. Jemielity S, Wang JJ, Chan YK, Ahmed AA, Li W, Monahan S, Bu X, Farzan M, Freeman GJ, Umetsu DT, Dekruyff RH, Choe H. 2013. TIM-family proteins promote infection of multiple enveloped viruses through virion-associated phosphatidylserine. PLoS Pathog 9:e1003232.

46. Richard AS, Zhang A, Park SJ, Farzan M, Zong M, Choe H. 2015. Virion-associated phosphatidylethanolamine promotes TIM1-mediated infection by Ebola, dengue, and West Nile viruses. Proc Natl Acad Sci U S A 112:14682–7.

47. Brouillette RB, Phillips EK, Patel R, Mahauad-Fernandez W, Moller-Tank S, Rogers KJ, Dillard JA, Cooney AL, Martinez-Sobrido L, Okeoma C, Maury W. 2018. TIM-1 Mediates Dystroglycan-Independent Entry of Lassa Virus. J Virol 92.

48. Zhang L, Richard AS, Jackson CB, Ojha A, Choe H. 2020. Phosphatidylethanolamine and Phosphatidylserine Synergize To Enhance GAS6/AXL-Mediated Virus Infection and Efferocytosis. J Virol 95.

49. Acciani MD, Lay Mendoza MF, Havranek KE, Duncan AM, Iyer H, Linn OL, Brindley MA. 2021. Ebola virus requires phosphatidylserine scrambling activity for efficient budding and optimal infectivity. J Virol doi:10.1128/JVI.01165-21:JVI0116521.

50. Dejarnac O, Hafirassou ML, Chazal M, Versapuech M, Gaillard J, Perera-Lecoin M, Umana-Diaz C, Bonnet-Madin L, Carnec X, Tinevez JY, Delaugerre C, Schwartz O, Roingeard P, Jouvenet N, Berlioz-Torrent C, Meertens L, Amara A. 2018. TIM-1 Ubiquitination Mediates Dengue Virus Entry. Cell Rep 23:1779–1793.

51. Amara A, Mercer J. 2015. Viral apoptotic mimicry. Nat Rev Microbiol 13:461–9.

52. Lay Mendoza MF, Acciani MD, Levit CN, Santa Maria C, Brindley MA. 2020. Monitoring Viral Entry in Real-Time Using a Luciferase Recombinant Vesicular Stomatitis Virus Producing SARS-CoV-2, EBOV, LASV, CHIKV, and VSV Glycoproteins. Viruses 12.

53. Cao W, Henry MD, Borrow P, Yamada H, Elder JH, Ravkov EV, Nichol ST, Compans RW, Campbell KP, Oldstone MB. 1998. Identification of alpha-dystroglycan as a receptor for lymphocytic choriomeningitis virus and Lassa fever virus. Science 282:2079–81.

54. Jae LT, Raaben M, Riemersma M, van Beusekom E, Blomen VA, Velds A, Kerkhoven RM, Carette JE, Topaloglu H, Meinecke P, Wessels MW, Lefeber DJ, Whelan SP, van Bokhoven H, Brummelkamp TR. 2013. Deciphering the glycosylome of dystroglycanopathies using haploid screens for lassa virus entry. Science 340:479–83.

55. Kunz S, Rojek JM, Kanagawa M, Spiropoulou CF, Barresi R, Campbell KP, Oldstone MB. 2005. Posttranslational modification of alpha-dystroglycan, the cellular receptor for arenaviruses, by the glycosyltransferase LARGE is critical for virus binding. J Virol 79:14282–96.

56. Krejbich-Trotot P, Denizot M, Hoarau JJ, Jaffar-Bandjee MC, Das T, Gasque P. 2011. Chikungunya virus mobilizes the apoptotic machinery to invade host cell defenses. FASEB J 25:314–25.

57. Shimojima M, Takada A, Ebihara H, Neumann G, Fujioka K, Irimura T, Jones S, Feldmann H, Kawaoka Y. 2006. Tyro3 family-mediated cell entry of Ebola and Marburg viruses. J Virol 80:10109–16.

58. Fast P. 1964. Insect lipids: A review. Entomological Society of Canada, Ottawa.

59. Luukkonen A, Brummer-Korvenkontio M, Renkonen O. 1973. Lipids of cultured mosquito cells (Aedes albopictus). Comparison with cultured mammalian fibroblasts (BHK 21 cells). Biochim Biophys Acta 326:256–61.

60. Shiomi A, Nagao K, Yokota N, Tsuchiya M, Kato U, Juni N, Hara Y, Mori MX, Mori Y, Ui-Tei K, Murate M, Kobayashi T, Nishino Y, Miyazawa A, Yamamoto A, Suzuki R, Kaufmann S, Tanaka M, Tatsumi K, Nakabe K, Shintaku H, Yesylevsky S, Bogdanov M, Umeda M. 2021. Extreme deformability of insect cell membranes is governed by phospholipid scrambling. Cell Rep 35:109219.

61. Nanbo A, Maruyama J, Imai M, Ujie M, Fujioka Y, Nishide S, Takada A, Ohba Y, Kawaoka Y. 2018. Ebola virus requires a host scramblase for externalization of phosphatidylserine on the surface of viral particles. PLoS Pathog 14:e1006848.

62. Yin P, Kielian M. 2021. BHK-21 Cell Clones Differ in Chikungunya Virus Infection and MXRA8 Receptor Expression. Viruses 13.

63. Basore K, Kim AS, Nelson CA, Zhang R, Smith BK, Uranga C, Vang L, Cheng M, Gross ML, Smith J, Diamond MS, Fremont DH. 2019. Cryo-EM Structure of Chikungunya Virus in Complex with the Mxra8 Receptor. Cell 177:1725–1737 e16.

64. Song H, Zhao Z, Chai Y, Jin X, Li C, Yuan F, Liu S, Gao Z, Wang H, Song J, Vazquez L, Zhang Y, Tan S, Morel CM, Yan J, Shi Y, Qi J, Gao F, Gao GF. 2019. Molecular Basis of Arthritogenic Alphavirus Receptor MXRA8 Binding to Chikungunya Virus Envelope Protein. Cell 177:1714–1724 e12.

65. Silva LA, Khomandiak S, Ashbrook AW, Weller R, Heise MT, Morrison TE, Dermody TS. 2014. A single-amino-acid polymorphism in Chikungunya virus E2 glycoprotein influences glycosaminoglycan utilization. J Virol 88:2385–97.

66. Ashbrook AW, Burrack KS, Silva LA, Montgomery SA, Heise MT, Morrison TE, Dermody TS. 2014. Residue 82 of the Chikungunya virus E2 attachment protein modulates viral dissemination and arthritis in mice. J Virol 88:12180–92.

67. Hawman DW, Fox JM, Ashbrook AW, May NA, Schroeder KMS, Torres RM, Crowe JE, Jr., Dermody TS, Diamond MS, Morrison TE. 2016. Pathogenic Chikungunya Virus Evades B Cell Responses to Establish Persistence. Cell Rep 16:1326–1338.

68. Bernard E, Hamel R, Neyret A, Ekchariyawat P, Moles JP, Simmons G, Chazal N, Despres P, Misse D, Briant L. 2015. Human keratinocytes restrict chikungunya virus replication at a post-fusion step. Virology 476:1–10.

69. Bauer T, Zagorska A, Jurkin J, Yasmin N, Koffel R, Richter S, Gesslbauer B, Lemke G, Strobl H. 2012. Identification of Axl as a downstream effector of TGF-beta1 during Langerhans cell differentiation and epidermal homeostasis. J Exp Med 209:2033–47.

70. Freeman GJ, Casasnovas JM, Umetsu DT, DeKruyff RH. 2010. TIM genes: a family of cell surface phosphatidylserine receptors that regulate innate and adaptive immunity. Immunol Rev 235:172–89.

71. Wang Q, Imamura R, Motani K, Kushiyama H, Nagata S, Suda T. 2013. Pyroptotic cells externalize eat-me and release find-me signals and are efficiently engulfed by macrophages. Int Immunol 25:363–72.

72. Pietkiewicz S, Schmidt JH, Lavrik IN. 2015. Quantification of apoptosis and necroptosis at the single cell level by a combination of Imaging Flow Cytometry with classical Annexin V/propidium iodide staining. J Immunol Methods 423:99–103.

73. Sourisseau M, Schilte C, Casartelli N, Trouillet C, Guivel-Benhassine F, Rudnicka D, Sol-Foulon N, Le Roux K, Prevost MC, Fsihi H, Frenkiel MP, Blanchet F, Afonso PV, Ceccaldi PE, Ozden S, Gessain A, Schuffenecker I, Verhasselt B, Zamborlini A, Saib A, Rey FA, Arenzana-Seisdedos F, Despres P, Michault A, Albert ML, Schwartz O. 2007. Characterization of reemerging chikungunya virus. PLoS Pathog 3:e89.

74. Young AR, Locke MC, Cook LE, Hiller BE, Zhang R, Hedberg ML, Monte KJ, Veis DJ, Diamond MS, Lenschow DJ. 2019. Dermal and muscle fibroblasts and skeletal myofibers survive chikungunya virus infection and harbor persistent RNA. PLoS Pathog 15:e1007993.

75. Lentscher AJ, McCarthy MK, May NA, Davenport BJ, Montgomery SA, Raghunathan K, McAllister N, Silva LA, Morrison TE, Dermody TS. 2020. Chikungunya virus replication in skeletal muscle cells is required for disease development. J Clin Invest 130:1466–1478.

76. Moser LA, Boylan BT, Moreira FR, Myers LJ, Svenson EL, Fedorova NB, Pickett BE, Bernard KA. 2018. Growth and adaptation of Zika virus in mammalian and mosquito cells. PLoS Negl Trop Dis 12:e0006880.

77. Reyes-Ruiz JM, Osuna-Ramos JF, Bautista-Carbajal P, Jaworski E, Soto-Acosta R, Cervantes-Salazar M, Angel-Ambrocio AH, Castillo-Munguia JP, Chavez-Munguia B, De Nova-Ocampo M, Routh A, Del Angel RM, Salas-Benito JS. 2019. Mosquito cells persistently infected with dengue virus produce viral particles with host-dependent replication. Virology 531:1–18.

78. Hafer A, Whittlesey R, Brown DT, Hernandez R. 2009. Differential incorporation of cholesterol by Sindbis virus grown in mammalian or insect cells. J Virol 83:9113–21.

79. Carro AC, Damonte EB. 2013. Requirement of cholesterol in the viral envelope for dengue virus infection. Virus Res 174:78–87.

80. Hanna SL, Pierson TC, Sanchez MD, Ahmed AA, Murtadha MM, Doms RW. 2005. N-linked glycosylation of west nile virus envelope proteins influences particle assembly and infectivity. J Virol 79:13262–74.

81. Lim PY, Louie KL, Styer LM, Shi PY, Bernard KA. 2010. Viral pathogenesis in mice is similar for West Nile virus derived from mosquito and mammalian cells. Virology 400:93–103.

82. Boylan BT, Moreira FR, Carlson TW, Bernard KA. 2017. Mosquito cell-derived West Nile virus replicon particles mimic arbovirus inoculum and have reduced spread in mice. PLoS Negl Trop Dis 11:e0005394.

83. Acharya D, Paul AM, Anderson JF, Huang F, Bai F. 2015. Loss of Glycosaminoglycan Receptor Binding after Mosquito Cell Passage Reduces Chikungunya Virus Infectivity. PLoS Negl Trop Dis 9:e0004139.

84. Dunbar CA, Rayaprolu V, Wang JC, Brown CJ, Leishman E, Jones-Burrage S, Trinidad JC, Bradshaw HB, Clemmer DE, Mukhopadhyay S, Jarrold MF. 2019. Dissecting the Components of Sindbis Virus from Arthropod and Vertebrate Hosts: Implications for Infectivity Differences. ACS Infect Dis 5:892–902.

85. van den Eijnde SM, Boshart L, Baehrecke EH, De Zeeuw CI, Reutelingsperger CP, Vermeij-Keers C. 1998. Cell surface exposure of phosphatidylserine during apoptosis is phylogenetically conserved. Apoptosis 3:9–16.

86. Ran FA, Hsu PD, Wright J, Agarwala V, Scott DA, Zhang F. 2013. Genome engineering using the CRISPR-Cas9 system. Nat Protoc 8:2281–2308.

87. Ramakrishnan MA. 2016. Determination of 50% endpoint titer using a simple formula. World J Virol 5:85–6.

88. McCarthy MK, Davenport BJ, Reynoso GV, Lucas ED, May NA, Elmore SA, Tamburini BA, Hickman HD, Morrison TE. 2018. Chikungunya virus impairs draining lymph node function by inhibiting HEV-mediated lymphocyte recruitment. JCI Insight 3.

